# The impact of various microplastics on bacterial community and antimicrobial resistance genes in Norwegian and South African wastewater

**DOI:** 10.1101/2024.05.22.595281

**Authors:** Tam Thanh Tran, Kabelo Stephans Stenger, Marte Strømmen, Cornelius Carlos Bezuidenhout, Odd-Gunnar Wikmark

## Abstract

Wastewater treatment plants (WWTPs) may serve as hotspots for pathogens and promote antimicrobial resistance (AMR). Plastic debris in wastewater could further contribute to AMR dissemination. The aim of this study is to investigate the impact of various microplastic types on bacterial communities and AMR gene abundance in Norwegian and South African wastewater. Microcosm experiments were designed as follows: Five manufactured microplastic pellet types were used for testing and two rock aggregate types were used as controls. In addition, each material type was subjected to artificial aging treatments using either ultra-violet light or hydrogen peroxide. Each material was incubated in flasks containing inlet/outlet wastewater obtained from Norwegian/South African WWTPs. Nucleic acids were extracted after a one-week incubation period. The detection of the *bla*_FOX_ *and bla*_MOX_ genes was performed using quantitative PCR. Extracted DNA was sequenced using a MinION device. Non-metric multi-dimensional scaling plot on full-length 16S sequencing data at the species level showed samples were clustered into distinct material groups. These results were in line with the ANOSIM test showing significant p-values in both Norwegian and South African WWTP settings. Indicator species analysis showed a strong association between many Acinetobacter species with the plastic group than the rock group. Aging treatment using hydrogen peroxide showed some effects on microbial. The abundance of *bla*_FOX_ and *bla*_MOX_ genes in the Norwegian wastewater outlet were generally lower compared to those in the inlet, though results were contrary in South African wastewater samples. The relative abundance of AMR genes seemed to be increased on several plastic types (PET, PE, PLA) but decreased on PVC-A. WWTP treatments in this study did not effectively reduce the abundance of AMR genes. In addition, plastic categories were shown to play a pivotal role in developing distinct bacterial communities and AMR profiles.

## 1. Introduction

Antimicrobial resistance (AMR) is one of top 10 global threats according to the World Health Organization WHO (Nations, 2023). Deaths attributable to AMR continue to rise alarmingly towards the number of 10 million deaths annually which was predicted in the 2016 report (O’Neill, 2016). In 2019, death toll associated with AMR was reported to be about 4.95 million which exceeded other death causes such HIV/AIDS, breast cancer or malaria (Collaborators, 2022). The Quadripartite organizations have developed ‘One Health Priority Research Agenda’ to mitigate this threat and support national action plan implementation and achievement of the Sustainable Development Goals for 2030 (Organization et al., 2023). One Health is a term to describe the health-related interconnection between human, animal and their living environments (Atlas, 2013; Prata et al., 2022). The One Health approach has been widely adopted by many organizations/government bodies such as EU, WHO and the United Nations (Commission, 2017; Organization et al., 2023; Programme, 2023).

Human and animal effluents and solid wastes from humans (including healthcare facilities and community settings) and from agri-food systems (including consumers) have been known as sources of pathogens and AMR genes (Sambaza and Naicker, 2023). Waste streams excreted from anthropogenic activities largely contributes to the dissemination of pathogens and AMR genes into the surrounding environments if they do not get treated properly, which mainly takes place at wastewater treatment plants (WWTPs) (Honda et al., 2023; Sambaza and Naicker, 2023). However, not all treatment methods result in a safe discharge of the effluent into the environment.

There are different treatment processes that can be used either alone or in combination to treat wastewater such as membrane filtration, coagulation, adsorption, and advanced oxidation processes (Sambaza and Naicker, 2023). Depending on geographical locations, wastewater profiles are drastically different, hence same treatment method could lead to various outcomes in wastewater discharge. Selection of appropriate treatment processes for a specific location is critical in eliminating/reducing the disposal of pollutants into the environment as well as in AMR monitoring. WWTPs processes are reported to remove over 90% of microplastics in the final effluent (Akarsu et al., 2020; Alavian Petroody et al., 2020; Blair et al., 2019), with the smallest particles showing the lowest removal efficiency (Alavian Petroody et al., 2020). For instance, WWTPs in China (Long et al., 2019; Zhang et al., 2021), UK (Blair et al., 2019), and Turkey (Akarsu et al., 2020) reported daily microplastics discharge of approximately 2.87 × 10^8^ - 6.5 × 10^8^ microplastics, 1.2 x 10^7^ microplastics, and 1.8 × 10^7^ microplastics, respectively. These WWTPs effectively eliminated microplastics with removal efficiency of 96% - 97%, 96% and 97% respectively. Despite the high removal efficiency large amounts of microplastics escape from WWTPs into aquatic environments, as such WWTP’s remain a major emission source of microplastics (Akarsu et al., 2020).

WWTP are collection points for microplastics present in domestic wastewater, industrial wastewater, and rainwater runoff. Therefore, microplastics present in WWTP could pose a risk in aquatic environments owing to their ability to act as vectors of other environmental contaminants (antibiotic, ARGs, and pathogens) present in wastewater. For instance, microplastics can act as carriers for pathogens derived from wastewater such as *Arcobacter* spp., *Aeromonas* spp., *and Vibrio* spp.(Lai et al., 2022), while enhancing horizontal gene transfer of ARGs in wastewater (Cheng et al., 2022). Moreover, microplastics can alter the distribution and removal of microbial communities and ARGs in treated wastewater (Eckert et al., 2018; Galafassi et al., 2021; Yang et al., 2022). In view of these factors, microplastics discharged in treated effluent serves as an important pathway for propagating antibiotic resistance in both terrestrial and aquatic food chains, posing a health threat to both human and livestock (Lai et al., 2022).

The main purpose of this study was to investigate the impact of various microplastic types on bacterial community and AMR gene abundance in both Norwegian and South African WWTPs. The study also further investigated the effect of two artificial plastic aging treatments on these bacterial communities.

## 2. Material and Method

### 2.1 Experiment set-up

Inlet and outlet wastewater samples were obtained from a local wastewater treatment facility in Tromsø, Norway and Potchefstroom, South Africa. A volume of 100ml wastewater was added into sterile flasks containing each material type (plastics or rocks) with an amount equivalent to 10ml volume. A sterile flask containing only wastewater was also included at the same time. These flasks were incubated at room temperature for seven days in a static condition.

Seven plastic types in pellet forms were used in this study: High-density polyethylene (HPDE), Polyvinyl chloride (PVC.A), Polyethylene terephthalate (PET), Polylactic acid (PLA), Polyethylene (PE) (purchased from Plastics SA, South African). Two types of rocks were purchased from a local store in Tromsø (Pets Tromsø Dyrebutikk): Black rock (Merkur 3-5mm, 3Ltr, AT20503), White rock (Sirius – Hvid AT22410 3-5 MM).

Before being used, above materials were sterilized by being soaked in bleach for 30 minutes, then transferred to ethanol 70% for 5 minutes and left air-dry in the biosafety cabinet. Aged materials were treated with either UV light with a wavelength of 254nm or hydrogen peroxide 33% (Catalog no. 23613.297, VWR Chemicals BDH®).

### 2.2 DNA extraction and quantification

DNA was extracted using phenol-chloroform protocol (Sambrook and Russell, 2006). Specifically, plastics/rocks were collected and washed once with 10ml of phosphate-buffered saline (PBS) + tween 0.1% buffer. Then 10ml of the same buffer was added into plastic pellets and vortex low to high speed for 1min to resuspend bacterial biofilm from plastics/rocks. Cells were harvested by being centrifuged at 12,000g, 1min (repeating the steps until finished), and the liquid was discarded. The cell pellet was resuspended in 300 µl PBS + tween 0.1% buffer; then 97µl lysozyme solution (10mg/ml) and 1 ul RNAse A was added and incubated at 37°C for 30mins. Subsequently lysis solution which was composed of 40µl of 25% SDS, 1ul Proteinase K and 100 µl of 5 M NaCl was added to lyse the cells at 65°C for 30 min. The protein was precipitated out by the addition of 500µl of 7.5M ammonium acetate and leaving on ice for 20 mins. Proteins were then removed by centrifuging at 20,000 g for 10 mins at 4°C. Aqueous solution was then transferred to a new tube containing 800µl phenol:chloroform:isoamyl alcohol and inverted several times before being centrifuged at 20,000g for 10mins at 4°C. The top phase was transferred to a new tube containing 800 µl chloroform. Inverting and centrifuging steps were repeated as above before the top phase was transferred to a new tube containing 1ml of isopropanol and incubated on ice for 30 mins to precipitate DNA. DNA pellet was collected by centrifuging at 20,000g, 10mins at 4°C, washed twice with 1ml of 70% ethanol dissolved in 100 µl of Tris buffer.

DNA was quantified using both NanoDrop 2000c spectrophotometer (ThermoFisher Scientific, Oslo, Norway) and Qubit 4 Fluorometer (ThermoFisher Scientific, Oslo, Norway) according to the manufacturer’s protocol.

### 2.3 Quantitative PCR analysis to detect and quantify AmpC β-lactamase gene groups

Quantification of AmpC β-lactamase gene groups (*bla*_FOX_ and *bla*_MOX_/ *bla*_CMY_) was performed using QuantStudioTM 3 System v1.4.3 (Applied Biosystems, Thermo Fisher Scientific, USA). The reaction mixture consisted of 1x TaqManTM Fast Advanced MasterMix (Thermo Fisher Scientific, USA), 1x TaqMan assay, DNA template, and nuclease-free water to a total reaction volume of 10 µl. Thermal cycling conditions consisted of a holding stage with step 1 at 50°C for 2 minutes and step 2 at 95°C for 10 minutes. The PCR stage consisted of 40 cycles with step 1 at 95°C for 15 seconds followed by a final step 2 at 60°C for 1 minute.

The following FAM fluorescent dyes were used for quantification: Pa04646126 (*bla*_FOX_) and Pa04646156_s1 (*bla*_MOX_/ *bla*_CMY_). Standard curves were generated using positive control samples for each target gene containing known copies using a ten-fold dilutions series in triplicates ranging between 20 000 to 2 copies. Amplification efficiencies of standard curves between 90% and 110% (E = 10−1/slope − 1) and R2 > 0.97 were deemed as reliable (Coertze and Bezuidenhout, 2020).

### 2.4 MinION sequencing protocol

DNA sequencing was performed on either a flongle FLO-FLG001 or MinION FLO-MIN106 flow cell (R9.4.1) using long-read Oxford Nanopore MinION platform (Oxford Nanopore Technologies Ltd., Oxford, UK). DNA libraries were prepared and loaded on flow cells following the default manufacturer protocol for 16S barcoding (the SQK-RAB204 or SQK-16S024 kit) or rapid barcoding (the SQK-RBK004 kit) for metagenomic sequencing. MinION Mk1C installed with software version 23.04.5 (Oxford Nanopore Technologies) was used to perform sequencing, demultiplexing, base-calling, and quality filtering of the raw reads.

### 2.5 Bioinformatic analyses

Full-length 16S fastq reads were further analysed using Fastq 16S v2022.01.07 on EPI2ME Desktop Agent v3.5.5 with default settings: min qscore of 7, BLAST e-value filter of 0.01, minimum coverage of 30%, min identity of 77%. The reports were then exported and saved as csv files so that further analyses were performed using R or python programming.

Metagenomic fastq reads were analyzed directly using Fastq Antimicrobial Resistance v2023.04.26 EPI2ME Desktop Agent v. 3.5.5 with default settings. Results were exported in tsv files in the epi2me output folder, and further analysis was performed using python programming. Metagenomic fastq reads were further assembled using the Flye v. 2.9.2 tool (Kolmogorov et al., 2020). The assemblies were used as the input to MOB-suite v. 3.1.0 to characterize the plasmid content, and StarAMR v. 0.9.1 to detect antibiotic resistance genes based on the resfinder resistance gene database (Bharat et al., 2022; Robertson and Nash, 2018; Zankari et al., 2012).

All sequencing data are deposited in the National Center for Biotechnology Information (NCBI) Sequence Read Archive under BioProject ID PRJNA1061070.

### 2.6 Visualization and statistical analyses

Statistical analysis was performed using either JASP software (Version 0.17.1) or R version 4.2.2 (2022-10-31 ucrt) on RStudio 2023.06.0 Build 421 (© 2009-2023 Posit Software, PBC). If assumptions for normality (Shapiro-Wilk) and homogeneity of variance (Levene’s Test) were met, t-test or one-way ANOVA was used to determine whether the difference was statistically significant (p < 0.05). If above assumption were not met, Mann-Whitney or Kruskal-Wallis Rank Sum Test was used instead. To determine whether the difference observed in the abundance of ARGs on microplastics substrates and natural substrates was statistically significant (p < 0.05), one-way ANOVA and Tukey’s HSD test were used.

For visualization, two R packages were used in this study, ‘vegan’ and ‘ggplot2’, to make Non-metric Multi-dimensional Scaling (NMDS) and bar plots. For statistical analyses, the non-parametric ANOSIM test with the number of permutations of 999 and the Bray–Curtis dissimilarity indices in the package ‘vegan’ was performed to determine if there is a statistical difference between the microbial communities of two or more groups of samples. The Indicator Species Analysis in the ‘indicspecies’ package was used to identify microbial species that are found more often in one treatment group compared to another.

## 3. Results

### 3.1 The impact of material and aging treatment on DNA extraction

DNA was extracted most from HPDE and PVC.A materials in the inlet wastewater set-up, while Black rock and PVC.A produced more DNA in the outlet wastewater set-up (Figure 1A). White rock produced the least DNA in both inlet and outlet wastewater set-ups. Given that assumptions were met, one-way ANOVA showed statistically significant p-values when comparing DNA concentration obtained from different materials in both set-ups (Inlet: *p* = 1.44−10^-6^; Outlet: *p* = 4.97−10^-4^). The Tukey’s HSD post-hoc test for pairwise comparisons showed these following group pairs significantly different in both set-ups (*p* < 0.05): HPDE – Black rock, PVC.A – PET, White rock – PVC.A.

**Figure 1:**
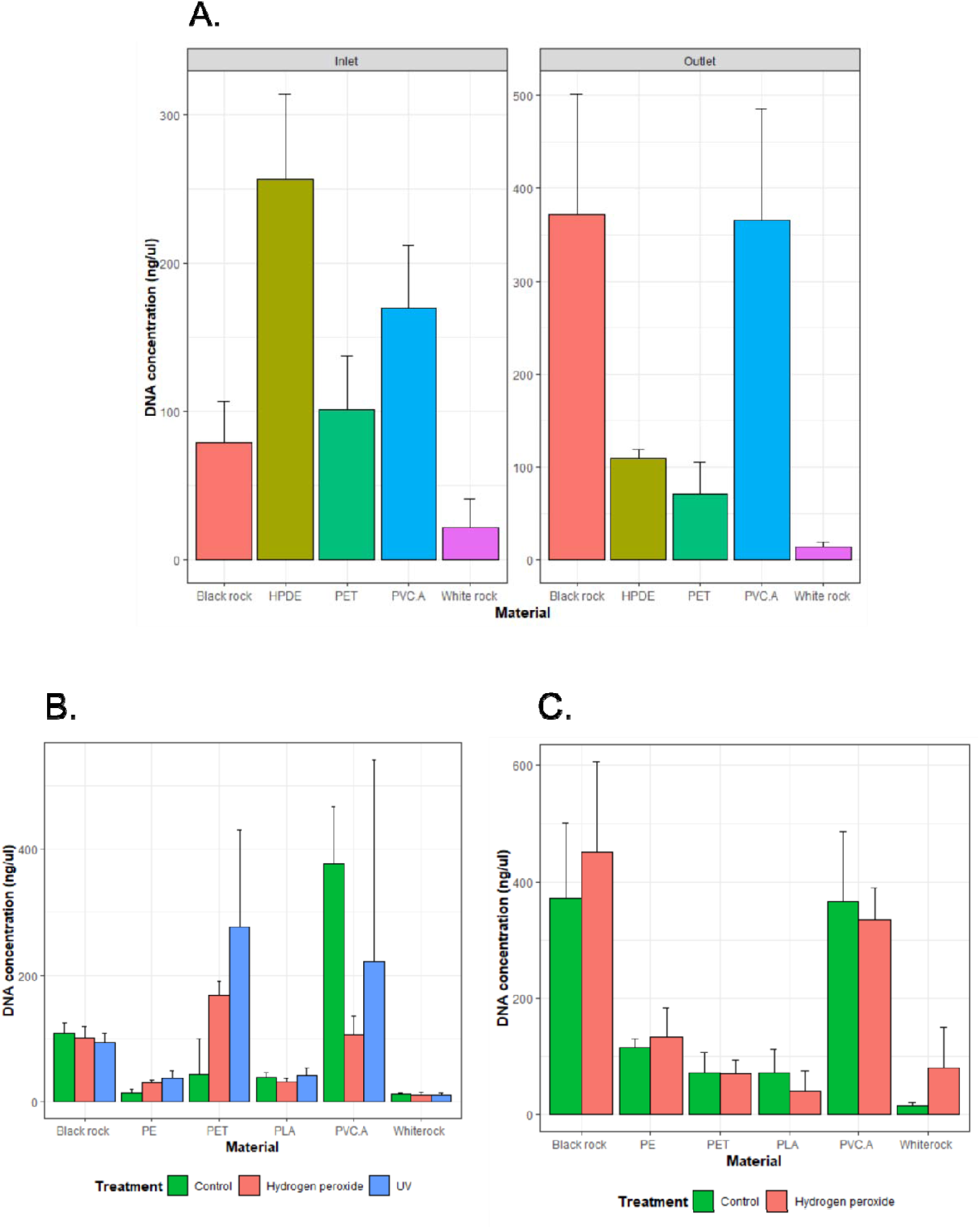
DNA obtained from A. Performing incubation with five different pristine materials: Black rock, HPDE, PET, PVC.A and White rock in Norwegian inlet and outlet wastewater set-up, B. Testing two aging treatments (Hydrogen peroxide or UV for 24-hour period) on six different materials: Black rock, PE, PET, PLA, PVC.A and White rock in Norwegian inlet wastewater, C. Testing one aging treatment (Hydrogen peroxide for 24-hour period) on six different materials: Black rock, PE, PET, PLA, PVC.A and White rock in Norwegian outlet wastewater.

Aging treatment and its duration posed different effects on extracted DNA depending on the material. For the inlet wastewater set-up, DNA concentrations appeared to increase significantly in aged PE (*p =* 0.01 using Hydrogen Peroxide and *p =* 0.02 using UV) and PET (*p =* 0.012 using Hydrogen Peroxide and *p =* 0.035 using UV) (Figure 1B). However, DNA concentration decreased in aged PVC.A after 24-hour treatment with hydrogen peroxide (*p =* 0.004). Shortening treatment period significantly increased DNA concentration extracted from aged PVC.A groups compared to control group (*p =* 0.02 using hydrogen peroxide and *p =* 0.026 using UV). Kruskal-Wallis’s test on all these samples showed a significant *p*-value if the grouping is based on material (*p =* 9.253−10^-6^) and insignificant *p*-value if the grouping is based on aging method (*p =* 0.81). For the outlet wastewater set-up, there was no significant difference in DNA concentrations between the control and aged groups for each material (Figure 1C). In addition, Kruskal-Wallis’s test also showed a significant difference when samples were grouped into different material types (*p* = 3.46−10^-5^), and insignificant difference when samples were grouped into non-aged/aged groups (*p =* 0.8).

### 3.2 Microbial genera from inlet and outlet wastewater and their colonization on plastics or rocks

For microcosm experiment with Norwegian inlet wastewater, Acidovorax and Dechloromonas are the most two prevalent genera present in three testing materials (Figure 2): Black rock (8.2%(117/1,420) and 4.9%(70/1,420), respectively), HPDE (10.4%(163/1,561) and 5.3%(83/1,561), respectively), PET (8.6%(32,380/374,833) and 7.1%(26929/374,833), respectively). Comamonas genus was found most prevalent from PVC.A pellets (44%(172/390)), followed by Acidovovax genus (7.1%(28/390)). Wastewater inlet seemed to have an even distribution among detected genera. DNA obtained from incubation using white rock in inlet wastewater was very little, resulting in very poor sequencing data.

**Figure 2:**
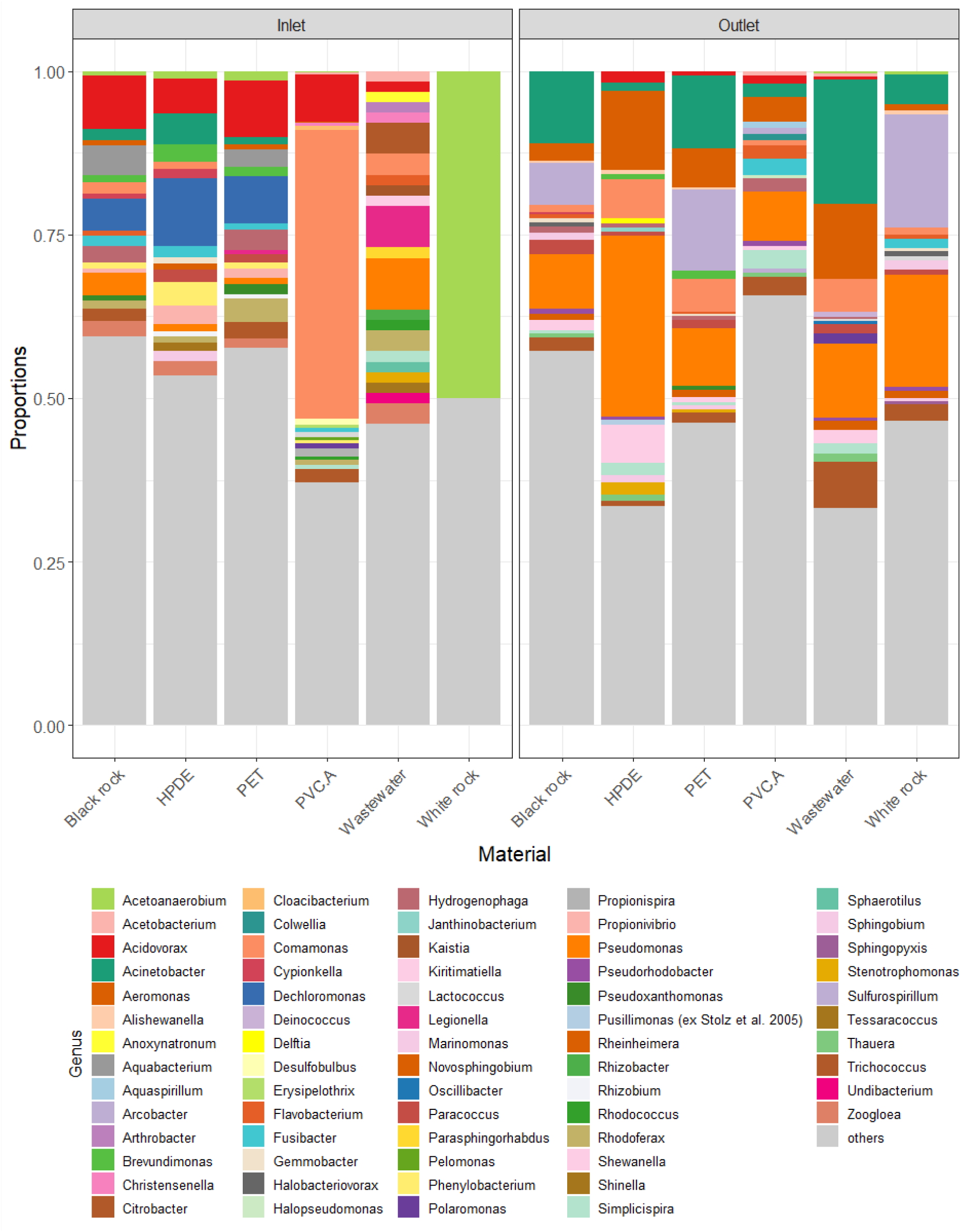
Top 20 bacterial genera detected in each testing material from Norwegian wastewater set-up.

For microcosm experiment with Norwegian outlet wastewater, Acinetobacter, Pseudomonas and Arcobacter were the most prevalent genera present in three testing materials (Figure 2): Black rock (11.1%(53,069/479,134), 8.3%(39,943/479,134) and 6.4%(31,031/479,134), respectively), White rock (4.6%(8,891/193,984), 17.1%(33,214/193,984) and 17.2%(33,380/193,984), respectively), PET (11.1%(58,934/533,037), 8.8%(46,730/533,037) and 12.3%(65,637/533,037), respectively). On the other hand, Pseudomonas and Aeromonas were found more on HPDE (27.6%(34,029/123,170) and 12.2%(15,004/123,170), respectively) and PVC.A (7.5%(35,236/465,346) and 3.8%(17,782/465,346), respectively). The top three genera found in outlet wastewater were Acinetobacter (19%(14,321/75,302)), Aeromonas (11.5%(8,662/75,302)) and Pseudomonas (11.3(8510/75,302)).

For microcosm experiment with South African inlet wastewater, Comamonas is among the top three prevalent genera present in all testing material (Figure 3): Black rock (4.6%(272/5974)), PET (8.9%(3714/41867)), PLA (10.8%(2593/23915), PVC.A (15.2%(9861/64828)). Other two prevalent genera found on these materials are: Acetoanaerobium (Black rock – 5.1%(302/5974), PET – 5.4%(2275/41867), PVC.A – 7.8%(5042/64828)), Stenotrophomonas (Black rock – 4%(239/5974), PLA – 10.9%(2610/23915)), Acinetobacter (PET – 25.3%(10613/41867), PLA – 9.6%(2299/23915)), Proteiniclasticum (PVC.A – 15.2%(9861/64828)). The top three genera found in the inlet wastewater were Acinetobacter (1.8%(307/16,987)), Comamonas (1.2%(203/16,987)) and Exiguobacterium (0.9%(159/16,987)).

**Figure 3:**
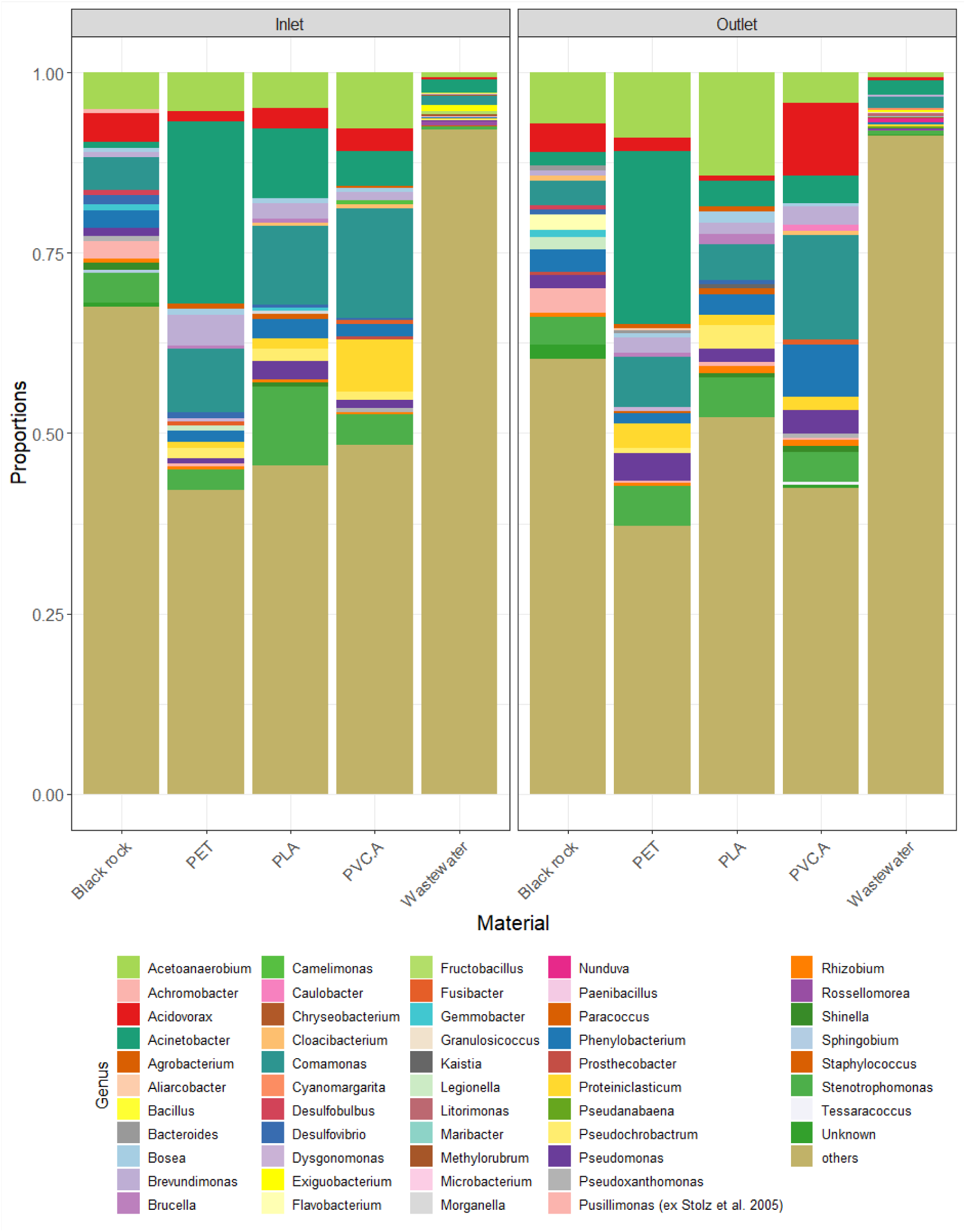
Top 20 bacterial genera detected in each testing material from South African wastewater set-up.

For microcosm experiment with South African outlet wastewater (Figure 3), following genera are among the top three prevalent genera present in testing materials: Comamonas (PET (7%(1,249/17,944)), PLA (5%(345/6,965), PVC.A (14.5%(2,619/17,948)); Acetoanaerobium (Black rock – 7.1%(1,268/17,932), PET – 9.1%(1,631/17,944), PLA – 14.3%(996/6,965)); Stenotrophomonas (Black rock – 3.7%(671/17,932), PLA – 5.5%(386/6,965)); Acinetobacter (PET – 24%(4,315/17,944)), Phenylobacterium (PVC.A – 7.2%(1,300/17,948)); Acidovorax (Black rock – 3.9%(708/17,932), PVC.A – 10.1%(1,812/17,948)). The top three genera found in outlet wastewater were Acinetobacter (2%(241/11,988)), Comamonas (1.7%(202/11,988)) and Acetoanaerobium (0.7%(84/11,988)).

### 3.3 The impact of material type on bacterial colonization at species level

The NMDS plots showed that samples were clustered well based on testing materials using bacterial abundance data as input (Figure 4A, 5A and 6A). This observation was consistent in both inlet and outlet wastewater set-ups, and for both Norwegian and South African samples.

**Figure 4:**
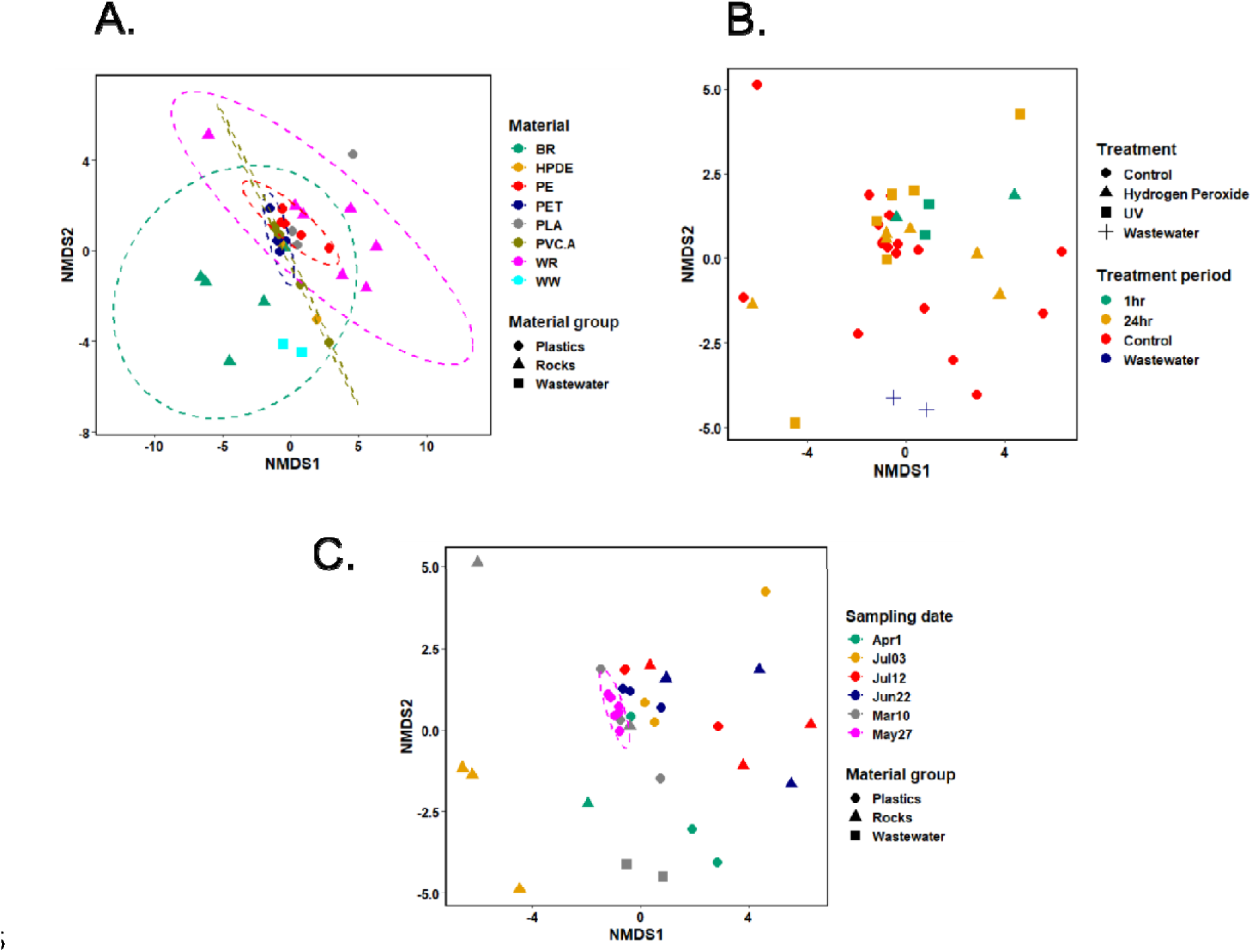
Non-metric Multi-dimensional Scaling (NMDS) analysis on bacterial community using full-length 16S rRNA gene dataset in microcosm experiment set-up by incubating of microplastics in Norwegian inlet wastewater. A) Seven different materials (BR: Black rock, WR: White rock, HPDE, PE, PET, PLA, PVC.A) and inlet wastewater (WW), B) Three treatment groups (Control, Hydrogen peroxide and UV) with either 1-hour or 24-hour periods and inlet wastewater (WW), C) Different sampling dates.

For Norwegian inlet wastewater set-up (Figure 4A), the ANOSIM test using Bray– Curtis’s dissimilarity matrices showed a very strong and statistically significant p-value of 5−10^-4^ with ANOSIM statistic R of 0.289 if samples were grouped into 3 categories (plastics, rocks or wastewater) and 8−10^-4^ with ANOSIM statistic R of 0.2501 if samples were grouped into specific types of rocks or plastics. When performing Indicator Species Analysis on three groups (plastics, rocks or wastewater), none of species were shown to be strongly associated with any of these groups. However, performing this analysis on specific materials showed a list of species that are likely to be associated with each material and their combined groups (Text S1). Interestingly, *Chryseobacterium treverense* showed some significant association with two plastic types (PET and PE).

For Norwegian outlet wastewater set-up (Figure 5A), the ANOSIM test using Bray– Curtis’s dissimilarity matrices also showed statistically significant p-value of 0.0087 with ANOSIM statistic R of 0.2662 if samples were grouped into 3 categories (plastics, rocks or wastewater) and 1−10^-4^ with ANOSIM statistic R of 0.2452 if samples were grouped into specific types of rocks or plastics. To our surprise, in this set-up, a list of species that had a strong association with plastic group compared to rock or wastewater only groups was able to be identified using Indicator Species Analysis (Text S2). Among those are nine Pseudomonas, four Stenotrophomonas, three Flavobacterium, and three Sphingobium species. In addition, performing Indicator Species Analysis on these data also listed a list of indicator species for each specific material and their combined groups (Text S3).

**Figure 5:**
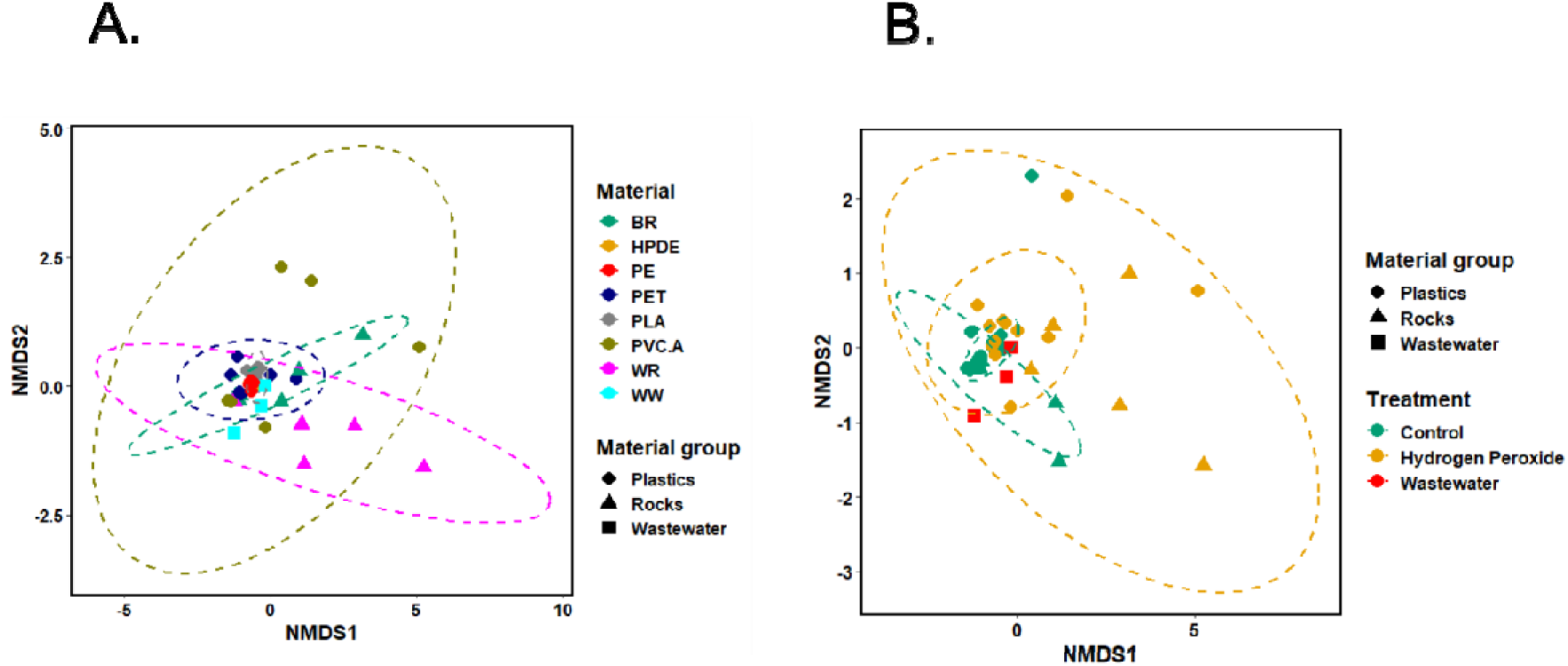
Non-metric Multi-dimensional Scaling (NMDS) analysis on bacterial community using full-length 16S rRNA gene dataset in microcosm experiment set-up by incubating of microplastics in Norwegian outlet wastewater. A) Seven different materials (BR: Black rock, WR: White rock, HPDE, PE, PET, PLA, PVC.A) and outlet wastewater (WW), B) Two treatment groups (Control, Hydrogen peroxide −24hrs) and outlet wastewater.

For South African wastewater set-up (Figure 6A), the ANOSIM test using Bray– Curtis’s dissimilarity matrices did not show statistically significant difference if samples were grouped into 2 categories: plastics and rocks (p-value of 0.06 with ANOSIM statistic R of 0.1379), but there was a significant difference if samples were grouped into specific types: BR, PET, PLA, PVC.A (1−10^-4^ with ANOSIM statistic R of 0.3268). When performing Indicator Species Analysis on two groups: plastics and rocks, 97 species were shown to be strongly associated with plastics and 368 species were strongly associated with rocks (only Black rock was used in this case) (Text S4). Notably, among those species, 38 Acinetobacter species showed significant association with the plastic group and none of Acinetobacter showed significant association with the rock group. Performing Indicator Species Analysis on specific materials showed a list of species that are likely to be associated with each material and their combined groups (Text S5). Interestingly, many Acinetobacter species (46 out of 86 species in total) showed a significant association in PET and PLA combined group.

**Figure 6:**
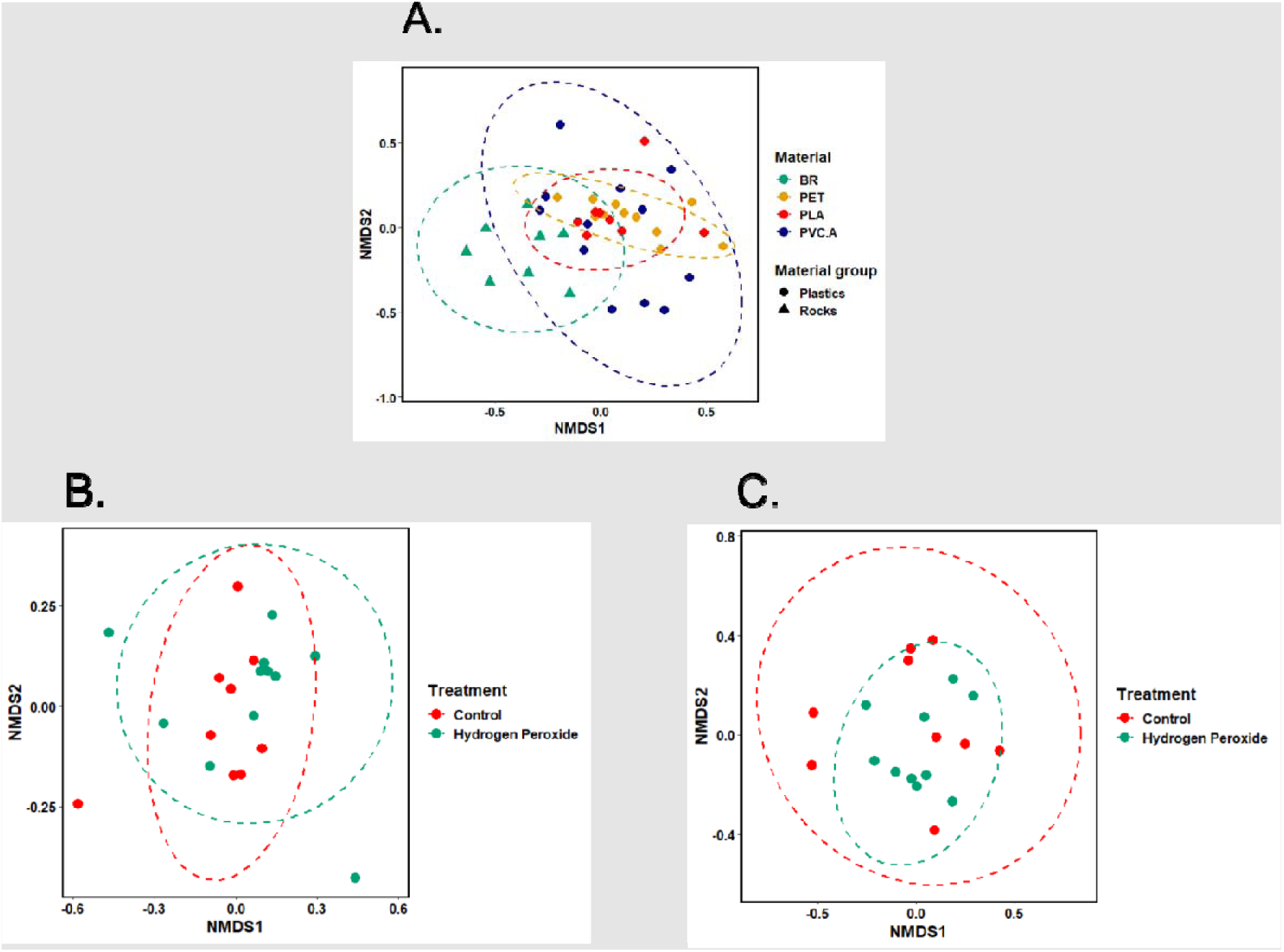
Non-metric Multi-dimensional Scaling (NMDS) analysis on bacterial community using full-length 16S rRNA gene dataset in microcosm experiment set-up by incubating of microplastics in South African wastewater. A) Four different materials (BR: Black rock, PET, PLA, PVC.A), B) Two treatment groups (Control and Hydrogen peroxide −24hrs) in inlet wastewater, C) Two treatment groups (Control and Hydrogen peroxide −24hrs) in outlet wastewater.

### 3.4 The impact of hydrogen peroxide and UV used in aging treatment on bacterial colonization at species level

For Norwegian inlet wastewater experiment set-up, samples from both treatment groups (Wastewater, Control, UV and Hydrogen peroxide) and treatment period groups (Wastewater, Control, 1-hour and 24-hour period) were scattered on NMDS plot (Figure 4B). There was also no significant difference between the microbial communities based on the type of treatment (ANOSIM statistic R: −0.04421 and p-value: 0.6942) or the period of treatment (ANOSIM statistic R: −0.04187 and p-value: 0.6824).

However, for Norwegian outlet wastewater experiment set-up, NMDS plot showed some clustering among different treatment groups (Wastewater, Control and Hydrogen peroxide) (Figure 5B). The ANOSIM test also showed a significant difference in microbial community among these groups (ANOSIM statistic R: 0.1431 and p-value: 0.0135). In outlet set-up, only hydrogen peroxide was used to treat materials for 24 hours.

Similar results were also observed for South African inlet and outlet wastewater set-ups (Figure 6B and 6C). There was no significant difference between aged and non-aged group in the inlet set-up (ANOSIM statistic R: 0.0101 and p-value: 0.3559), but significant difference in the outlet set-up (ANOSIM statistic R: 0.1868 and p-value: 0.0251). Materials were aged by being immersed in hydrogen peroxide for 24 hours.

### 3.5 Detection of AMR genes and mobile genetic sequences in Norwegian inlet and outlet wastewater using metagenomic approach

Many AMR genes were detected in both inlet and outlet wastewater set-ups using Epi2me pipeline on raw reads (Figure 7). The heatmap showed that certain AMR genes detected on several microplastics (PE, PET, PLA) abundantly increased in both inlet and outlet set-ups compared to those detected in wastewater or black rock samples. Those genes were found belonging to following functional groups: aminoglycoside resistance, efflux pump, elfamycin resistance, fluoroquinolone resistance and rifampicin resistance. To a lesser extent, the proliferation of AMR genes was also observed in beta-lactam resistance, sulfonamide resistance and tetracycline resistance. However, this was not the case for PVC.A on which AMR genes were detected less than those on black rock and wastewater samples. The relative abundances of total AMR genes for these samples were also in agreement with the above heatmap results in Figure 7.

**Figure 7:**
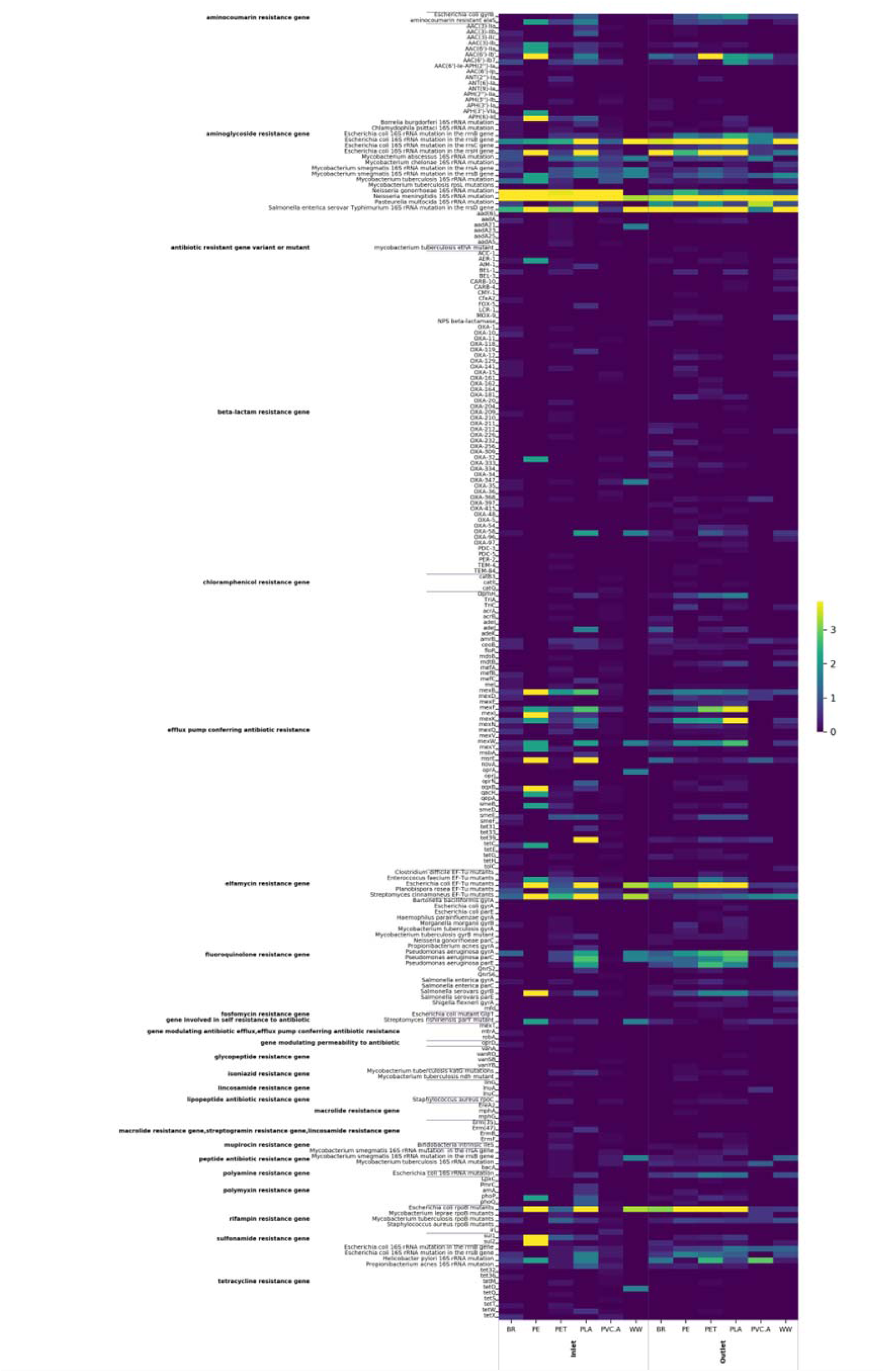
Heatmap diagram showing the relative abundance of AMR genes detected in both inlet and outlet wastewater set-ups.

Detection of AMR genes on assemblies also showed several acquired resistance genes (Table S1). Notably, these AMR genes were found only from plastic group (PLA, PET, PE). Many insertion sequences were detected on chromosomal assemblies from both inlet and outlet wastewater set-ups (Table S2). In the inlet wastewater, Tn3, IS6 and IS3 were the ones detected on PLA and PET groups. While in the outlet wastewater, a more variety of insertion sequences were detected, especially in PLA, PET and PE groups. Only two insertion sequences were detected on plasmid contigs from PET-outlet group (Table S3).

### 3.6 Quantification of *bla*_FOX_ and *bla*_MOX_ genes in inlet and outlet wastewater

In Norwegian samples, the *bla*_FOX_ and *bla*_MOX_ genes were generally found decreased in the outlet compared to those in the inlet wastewater set-up (Figure 8A and 8B). When comparing copy numbers of the *bla*_FOX_ gene in the outlet with those in the inlet wastewater set-up, significant decreases were found on aged PET (p=0.005) and aged PVC.A (p<0.001). When comparing copy numbers of the *bla*_MOX_ gene in the outlet wastewater with those in the inlet wastewater set-up, significant decreases were found on non-aged PVC.A (p<0.001) and aged PVC.A (p<0.001). When comparing between aged and non-aged materials, the inlet set-up showed that the *bla*_FOX_ gene copy numbers on aged-PET (p=0.045) and PVC.A (p=0.02) were significantly higher. Similarly, the *bla*_MOX_ gene copy number on aged PVC.A (p=0.04) was also significantly higher.

**Figure 8:** Gene copies per ng DNA of *bla*_FOX_ genes (A and C) and *bla*_MOX_ genes (B and D) inlet and outlet wastewater from Tromsø, Norway (A and B) and South African (C and D). **Treatment:** H-Hydrogen peroxide (aged); B-Bleach (control). **Significant:** P_1_ - between groups (inlet-outlet); P_2_ -between treatment (aged-control)

Meanwhile, in South African samples, the *bla*_FOX_ and *bla*_MOX_ genes were generally found increased in the outlet compared to those in the inlet wastewater set-up (Figure 8C and 8D). When comparing copy numbers of the *bla*_FOX_ genes in the outlet wastewater with those in the inlet wastewater set-up, significant increases were found in aged PET (p=0.006), aged PLA (p=0.013), aged Blackrock (p=0.003). When comparing copy numbers of the *bla*_MOX_ gene in the outlet wastewater with those in the inlet wastewater set-up, significant increase was found in aged PET (p<0.001). When comparing between aged and non-aged materials, the *bla*_FOX_ gene copy numbers in aged-PLA (p<0.001) and aged-Black rock (p<0.001) were significantly higher in the outlet set-up. The *bla*_MOX_ gene copy numbers were also significantly higher in aged PET (p=0.001) and aged PLA (p=0.006) in the inlet set-up, and in aged PET (p<0.001) in the outlet set-up.

## 4. Discussion

DNA amount extracted from different testing materials were shown to be significantly different among them. Very little amount of DNA was obtained from white rock, which was consistent across replicates between inlet and outlet set-ups. To our surprise, there were a great difference in extracted DNA between white rock and black rock, although they were purchased from the same pet store for the same use purpose as decorators in aquarium tank. We hypothesize that compositional difference is the main factor leading to different DNA amount extracted from these two rocks. The difference in total extracted DNA may be resulted from how well the materials can be served as a ‘habitat’ for microorganisms to grow. Information on original countries where these rocks were sourced were given by the production company Akvastabil, Eldorado A/S, Denmark. The white rock was from Sweden and the black rock was from Germany. They both were crushed after being excavated without any further treatment at the factory. The white rock, according to the company, also called ‘Dolomitkross’ (Dolomite crushes), is mainly composed of the mineral dolomite, CaMg(CO_3_)_2_ (King, 2005). It was shown that CaMg(CO_3_)_2_ has some antibacterial properties, and this explains poor DNA obtained from this white rock in both inlet and outlet set-ups (Han et al., 2022). There was also significant difference in extracted DNA among different types of testing microplastics. More surprisingly, the pattern was not consistent between inlet and outlet set-ups. For example, DNA obtained from black rock in the inlet set-up was less than that in the outlet set-up, while the opposite result happened with HPDE. An explanation could be the source of bacteria introduced to these materials. If a specific material favored the colonization of certain bacteria present in the inlet wastewater, the overall DNA outcome would then be better in the inlet set-up than the outlet set-up. This will also be further discussed as we investigate the effect of material on bacterial composition. DNA extracted here is the total environmental DNA which may include free DNA present in the wastewater samples. These free DNA could possibly be adsorbed on testing materials during the incubation (Hao et al., 2018; Wu et al., 2023). It was shown that different plastic types posed different adsorption affinity on environmental DNA (Wu et al., 2023).

There are mounting evidence that microplastics played an important role in the colonization of biofilm-forming microorganisms (Dudek et al., 2020; Gonzalez-Pleiter et al., 2021; Miao et al., 2019; Mishra et al., 2021). In our study, there was a significant effect of material on microbial community. This observation was seen in both inlet and outlet wastewater set-up and in both Norway and South African. The effect of material on microbial community was most obvious when samples were grouped in specific types of materials. This result is in line with several previous studies which showed a significant different in microbial community colonizing on different plastic types (Frere et al., 2018; Kirstein et al., 2018; Pinnell and Turner, 2019). However, results from a previous study suggested otherwise that plastic polymer type did not play a significant role in shaping biofilm communities (Dudek et al., 2020). Of note, there are a few differences in how to assess the role of plastic types on microbial communities between our study and theirs: microbial source, testing plastics, sequencing technology, taxonomic levels used as an input data and statistical models. Among those differences, we supposed that using different taxonomic levels as an input would be the main factor accounting for different outcomes. Because the effect of plastic types on bacterial community was also seen generally low in another study when looking at the genus level, however there were still some significant differences between biofilms on diverse polymer types (Kirstein et al., 2018). In our study, species was used as an input while genus was used in their study. Genus might not provide sufficient in-depth taxonomic resolution to determine the impact of plastic polymer type on bacterial community. In another study, it was shown that relative abundance in bacterial community at the genus level were significantly different between microplastics, surrounding water and rick microbial mats; however, it is unclear if this is also the case for different types of plastics (Gonzalez-Pleiter et al., 2021). The common observation that was seen across these studies was an increased richness in bacterial community in microplastics compared to those in water. In South African samples, bacteria from wastewater were clustered well and separately from the rest of bacteria present on material (data not shown).

Materials such as plastics or rocks exposed to chemicals or sunlight in nature are likely weathered through oxidation/photodegradation processes (Amelia et al., 2022; Andrady et al., 2022; Yousif and Haddad, 2013). Degradation mechanism when exposed to sunlight, also termed as ‘photodegradation’, has been well-studied and is caused by the absorption of photons from several wavelengths found in sunlight including infrared radiation, visible light, and ultraviolet light (Andrady et al., 2022; Yousif and Haddad, 2013). This mechanism can occur in the absence of oxygen (chain breaking or cross-linking) or the presence of oxygen (photooxidation). In this study, ultraviolet light was used to induce the photooxidative degradation process which may result in fragmentation and/or surface ablation (Andrady et al., 2022). Material exposed to hydrogen peroxide, a strong oxidizing agent, tended to be degraded or to have structural changes (Schrank et al., 2022). Recent studies mainly investigated the effect of aged microplastics on bacterial community from soil, river and coastal area. They showed that aged microplastics seemed to pose some effects on total biomass, metabolic pathway, enzyme activity and microbial community structure (Choi et al., 2021; Liu et al., 2022a; Shan et al., 2022). Notably, it was shown that aged microplastics significantly enriched potentially human pathogenic genes (Bao et al., 2022). Very few studies have been investigating the effect of aged microplastics on bacterial community in wastewater, although it showed that wastewater treatment plants changed morphology, sizes, and chemical properties of microplastics (Liu et al., 2022b). In our study, there was no significant effect of all aging treatments (UV and hydrogen peroxide with duration of 1 hour or 24 hours) on bacterial community from inlet wastewater. However, a significant effect on bacterial community from outlet wastewater was seen when materials were treated with hydrogen peroxide for 24 hours. There are a few explanations to be accounted for this difference: (1) these aging treatments were unlikely to degrade materials enough to create a significant shift in microbial community; (2) Samples were less diverse in outlet wastewater set-up; (3) the diversity of bacterial community was also likely reduced in outlet wastewater. According to a previous study, degradation on a molecular level was only observed for polyamide based on gel permeation chromatography analysis when hydrogen peroxide was used to treat different types of microplastics (Schrank et al., 2022). A study that showed the significant aging effect on microbial community had materials exposed to UV for an extended duration of at least 15 days (Bao et al., 2022). Also, it is important to point out that UVA was used to age materials instead of UVC in that study, and the aging treatment occurred at the same time as the incubation in that study.

Since COVID 19 pandemic, wastewater-based surveillance has been emerging as an alternative way to obtain data on population-wise infectious disease including AMR (Hendriksen et al., 2019; Lira et al., 2020; Sambaza and Naicker, 2023). Both the *bla*_FOX_ and *bla*_MOX_ genes, belong to AmpC-type β-lactamases, hydrolyse extended-spectrum cephalosporins and cephamycins (Ebmeyer et al., 2019). The qPCR results showed that both of *bla*_FOX_ and *bla*_MOX_ genes in the Norwegian wastewater outlet were generally lower compared to those in the inlet, while this seemed to be the opposite in South African wastewater samples. This could be due to the difference in wastewater treatment methods between these sites. In this Norwegian facility, a 300µM mesh filter was used to remove particles larger than 300µM from the inlet. Whereas, in South African treatment facility, conventional wastewater treatment is employed using mesh screens (3 - 12.5 mm). Although these genes were reduced in Norwegian wastewater outlet, the abundance of these genes were much higher in the Norwegian site in comparison with the South African site. This was because this Norwegian site was getting wastewater from the industrial area and a hospital nearby.

In recent years, metagenomic sequencing has been widely used to investigate the presence and abundance of AMR genes as well as mobile genetic elements in WWTPs (Che et al., 2019; Dai et al., 2022; Guo et al., 2017; Lira et al., 2020). Microorganisms were shown to form biofilm and colonize on microplastic particles. Therefore, microplastics can play a role of a carrier to facilitate horizontal transfer of AMR genes (Galafassi et al., 2021; Lai et al., 2022). To date, very few studies investigate the effect of microplastics on AMR gene abundance in WWTPs, even less so for specific plastic types. In our study, preliminary results showed that the proliferation of AMR genes were observed on most of materials in both inlet and outlet set-ups, especially more on certain microplastics. To our surprise, the abundance of AMR genes seems only increasing on certain plastic types (PE, PET and PLA), but decreasing in PVC.A compared to that on rocks or in wastewater samples. This observation was seen similar in both inlet and outlet set-ups. However, in another study, the abundance of AMR genes were higher on rocks and microplastics (PVC) compared to that in river water (Wu et al., 2019). In that study, only a single type of microplastics (PVC) was used for testing, and all the materials were incubated in the same bioreactor. Contrasting results regarding the effect of microplastics on potential pathogenic bacteria have been reported before in two independent studies (Galafassi et al., 2021; McCormick et al., 2014). In one study, microplastics supported the colonization of potentially pathogenic bacteria in comparison to the river-originated planktonic community (McCormick et al., 2014). While potentially pathogenic bacteria were found to be more abundant in the planktonic bacterial community in wastewater opposed to those on microplastics (Galafassi et al., 2021).

## 5. Conclusions

This study explored the role of different types of microplastics on bacterial community and AMR genes. It was shown that plastic type played a pivotal role in shaping bacterial community. Acinetobacter species showed a strong association with biofilm on plastic groups compared to that on rock groups. Aging treatment using ultra-violet did not show any significant effects on bacterial community profiles, while hydrogen peroxide treatment did. AMR genes were abundantly present in the outlet, indicating that WWTPs in this study were not effectively reduced the abundance of these genes.

To date, microplastics and antimicrobial resistance are two of the most important health-related anthropogenic pollution problems in the environment. Yet, very little practical data on the impact of microplastics on bacterial community and AMR genes were reported, even lesser for plastic types. Locating the source of these problems and building better management strategies are essential in mitigating AMR threat to humans, animals, and the environment.

## Acknowledgements

This study was supported by the Norwegian Research Council (MicroPlastResist— project number 288073 and MARMIB—project number 315812).

