## Supplement data for "The impact of various microplastics on bacterial community and antimicrobial resistance genes in Norwegian and South African wastewater"

Text S1: Microbial species that showed a statistically significant association in each material (Black rock, White rock, HPDE, PE, PET, PVC.A, PLA)/wastewater, and their combined groups using a specific R package (indicspecies) to perform Indicator Species Analysis with Norwegian inlet wastewater set-up.

```
Multilevel pattern analysis
 ---------------------------

 Association function: r.g
 Significance level (alpha): 0.05

 Total number of species: 5296
 Selected number of species: 42
 Number of species associated to 1 group: 41
 Number of species associated to 2 groups: 1
 Number of species associated to 3 groups: 0
 Number of species associated to 4 groups: 0
 Number of species associated to 5 groups: 0
 Number of species associated to 6 groups: 0
 Number of species associated to 7 groups: 0

 List of species associated to each combination:

 Group PE  #sps.  10
                                     stat p.value
Pseudomonas composti                0.816  0.0102 *
Rhizobium ipomoeae                  0.798  0.0180 *
Alishewanella longhuensis           0.775  0.0114 *
Leadbetterella byssophila DSM 17132 0.736  0.0141 *
Rheinheimera tangshanensis          0.707  0.0230 *
Aliterella antarctica               0.644  0.0381 *
Alishewanella solinquinati          0.642  0.0292 *
Taishania pollutisoli               0.642  0.0299 *
Alishewanella jeotgali KCTC 22429   0.622  0.0417 *
Stagnimonas aquatica                0.615  0.0305 *

 Group PET  #sps.  2
                            stat p.value
Rhizobium arsenicireducens 0.707  0.0199 *
Flavobacterium hercynium   0.682  0.0190 *

 Group PLA  #sps.  1
                         stat p.value
Exiguobacterium marinum 0.668  0.0217 *

 Group PVC.A  #sps.  28
                                      stat p.value
Marinobacter nauticus ATCC 49840     0.723  0.0208 *
Pseudomonas alkylphenolica           0.723  0.0208 *
Pseudomonas trivialis                0.708  0.0194 *
Pseudomonas rhodesiae                0.702  0.0208 *
Yersinia frederiksenii               0.699  0.0204 *
Aeromonas tecta                      0.694  0.0195 *
Janthinobacterium svalbardensis      0.693  0.0201 *
Pelosinus defluvii                   0.692  0.0223 *
Citrobacter freundii                 0.685  0.0235 *
Janthinobacterium aquaticum          0.683  0.0191 *
Pseudomonas grimontii                0.681  0.0227 *
Psychrosinus fermentans              0.681  0.0151 *
Yersinia kristensenii                0.678  0.0188 *
Pelosinus fermentans DSM 17108       0.670  0.0178 *
Aeromonas veronii                    0.669  0.0214 *
Oceanimonas doudoroffii              0.669  0.0255 *
Pseudomonas marginalis               0.667  0.0208 *
Pseudomonas asturiensis              0.648  0.0238 *
Pseudomonas meridiana                0.647  0.0386 *
Citrobacter werkmanii                0.639  0.0486 *
Anaerosinus glycerini                0.638  0.0176 *
Lelliottia amnigena                  0.637  0.0364 *
Pseudomonas veronii                  0.632  0.0307 *
Pseudomonas antarctica               0.629  0.0370 *
Yersinia aldovae                     0.625  0.0310 *
Desulfovibrio aerotolerans           0.615  0.0343 *
Aeromonas salmonicida subsp. smithia 0.613  0.0490 *
Aeromonas molluscorum                0.603  0.0496 *

 Group PE+PET  #sps.  1
                             stat p.value
Chryseobacterium treverense 0.636  0.0382 *
---
Signif. codes:  0 ‘***’ 0.001 ‘**’ 0.01 ‘*’ 0.05 ‘.’ 0.1 ‘ ’ 1
```

Text S2: Microbial species that showed a statistically significant association in each material group (plastics, rocks or wastewater) or their combined groups using a specific R package (indicspecies) to perform Indicator Species Analysis with Norwegian outlet wastewater set-up.

```
Multilevel pattern analysis
 ---------------------------

 Association function: r.g
 Significance level (alpha): 0.05

 Total number of species: 9455
 Selected number of species: 144
 Number of species associated to 1 group: 144
 Number of species associated to 2 groups: 0

 List of species associated to each combination:

 Group Plastics  #sps.  27
                                            stat p.value
Pseudomonas anguilliseptica                0.598  0.0176 *
Stenotrophomonas acidaminiphila            0.598  0.0151 *
Sphingobium naphthae                       0.567  0.0219 *
Pseudomonas borbori                        0.540  0.0277 *
Flavobacterium acidificum                  0.539  0.0282 *
Pseudomonas glareae                        0.539  0.0257 *
Shewanella oneidensis                      0.536  0.0264 *
Pseudomonas peli                           0.534  0.0454 *
Shewanella abyssi                          0.532  0.0308 *
Pseudomonas marincola                      0.525  0.0385 *
Thiomicrospira cyclica ALM1                0.523  0.0364 *
Pseudomonas helleri                        0.518  0.0496 *
Sphingobium xenophagum                     0.517  0.0364 *
Pseudomonas chlororaphis subsp. aurantiaca 0.516  0.0361 *
Xanthomonas translucens                    0.515  0.0393 *
Leucothrix mucor DSM 2157                  0.513  0.0330 *
Stenotrophomonas ginsengisoli              0.512  0.0428 *
Pseudomonas extremorientalis               0.511  0.0435 *
Pseudomonas composti                       0.505  0.0432 *
Flavobacterium kingsejongi                 0.504  0.0473 *
Stenotrophomonas maltophilia               0.504  0.0425 *
Sphingobium lucknowense F2                 0.500  0.0438 *
Sedimenticola selenatireducens             0.492  0.0476 *
Stenotrophomonas humi                      0.487  0.0362 *
Lysobacter caeni                           0.483  0.0458 *
Janthinobacterium aquaticum                0.464  0.0473 *
Flavobacterium lindanitolerans             0.446  0.0494 *

 Group Rocks  #sps.  15
                                stat p.value
Marinomonas aquiplantarum      0.548  0.0259 *
Winogradskyella thalassocola   0.516  0.0292 *
Marivita roseacus              0.478  0.0476 *
Psychrobacter aestuarii        0.476  0.0310 *
Halobacteriovorax litoralis    0.470  0.0455 *
Paracoccus alkenifer           0.470  0.0478 *
Rickettsia canadensis          0.463  0.0301 *
Aquimixticola soesokkakensis   0.463  0.0472 *
Steroidobacter agariperforans  0.462  0.0421 *
Cephaloticoccus primus         0.453  0.0461 *
Fulvimonas yonginensis         0.447  0.0262 *
Thalassotalea ponticola        0.447  0.0250 *
Jeotgalibacillus marinus       0.447  0.0263 *
Anoxybacillus amylolyticus     0.447  0.0272 *
Paracholeplasma brassicae 0502 0.441  0.0495 *

 Group Wastewater  #sps.  102
                                             stat p.value
Legionella quateirensis                     0.803  0.0030 **
Pelotomaculum thermopropionicum SI          0.765  0.0019 **
Listeria innocua                            0.756  0.0038 **
Rhodococcus rhodochrous                     0.721  0.0075 **
Novipirellula aureliae                      0.707  0.0038 **
Leptolinea tardivitalis                     0.707  0.0038 **
Yersinia nurmii                             0.702  0.0030 **
Friedmanniella luteola                      0.688  0.0101 *
Rhodococcus gannanensis                     0.688  0.0109 *
Spirosoma jeollabukense                     0.688  0.0115 *
Bacillus alkalitolerans                     0.685  0.0091 **
Auraticoccus cholistanensis                 0.674  0.0022 **
Nocardioides lianchengensis                 0.668  0.0070 **
Auraticoccus monumenti                      0.657  0.0121 *
Ornithinibacillus salinisoli                0.657  0.0128 *
Microlunatus sagamiharensis                 0.645  0.0003 ***
Microbacterium xylanilyticum                0.644  0.0189 *
Brevibacterium sanguinis                    0.644  0.0181 *
Rhodanobacter humi                          0.644  0.0189 *
Rarobacter faecitabidus                     0.643  0.0109 *
Microbacterium hydrothermale                0.619  0.0144 *
Peptoniphilus coxii                         0.617  0.0083 **
Roseomonas aquatica                         0.612  0.0074 **
Nitrosomonas europaea                       0.612  0.0050 **
Ilumatobacter fluminis YM22-133             0.604  0.0139 *
Shigella dysenteriae                        0.593  0.0187 *
Arthrobacter oryzae                         0.588  0.0206 *
Microbacterium mangrovi                     0.588  0.0339 *
Rhodopirellula heiligendammensis            0.584  0.0126 *
Pengzhenrongella sicca                      0.584  0.0144 *
Anaerofilum agile                           0.583  0.0046 **
Microbacterium profundi                     0.573  0.0142 *
Botrimarina mediterranea                    0.567  0.0272 *
Microbacterium lacus                        0.566  0.0074 **
Arthrobacter methylotrophus                 0.561  0.0020 **
Nocardioides alpinus                        0.556  0.0161 *
Brachybacterium avium                       0.556  0.0282 *
Microlunatus spumicola                      0.554  0.0024 **
Aureliella helgolandensis                   0.554  0.0213 *
Egibacter rhizosphaerae                     0.547  0.0345 *
Leucobacter chromiireducens                 0.547  0.0099 **
Agrococcus terreus                          0.545  0.0191 *
Pseudarthrobacter phenanthrenivorans Sphe3  0.545  0.0140 *
Flavimarina pacifica                        0.543  0.0252 *
Tetrasphaera australiensis                  0.540  0.0231 *
Serratia proteamaculans                     0.539  0.0192 *
Bacillus wudalianchiensis                   0.532  0.0397 *
Caloramator australicus RC3                 0.530  0.0170 *
Arthrobacter pascens                        0.529  0.0204 *
Caloramator indicus                         0.529  0.0175 *
Polaromonas glacialis                       0.522  0.0390 *
Diaminobutyricimonas aerilata               0.520  0.0276 *
Patulibacter minatonensis DSM 18081         0.519  0.0370 *
Demequina lutea                             0.515  0.0264 *
Desulfobulbus propionicus DSM 2032          0.510  0.0124 *
Serinibacter salmoneus                      0.506  0.0160 *
Gimesia maris                               0.503  0.0474 *
Desulforhopalus singaporensis               0.502  0.0168 *
Leucobacter japonicus                       0.501  0.0342 *
Clostridium estertheticum subsp. laramiense 0.500  0.0157 *
Fonticella tunisiensis                      0.499  0.0184 *
Desulfopila inferna                         0.496  0.0265 *
Arthrobacter psychrolactophilus             0.495  0.0495 *
Sedimentibacter acidaminivorans             0.494  0.0273 *
Advenella faeciporci                        0.494  0.0294 *
Litorihabitans aurantiacus                  0.491  0.0297 *
Sedimentibacter saalensis                   0.487  0.0316 *
Demequina oxidasica                         0.487  0.0234 *
Homoserinimonas aerilata                    0.479  0.0408 *
Christensenella timonensis                  0.472  0.0314 *
Microbacterium aurum                        0.471  0.0320 *
Paenibacillus medicaginis                   0.471  0.0274 *
Tessaracoccus massiliensis                  0.471  0.0308 *
Acutalibacter muris                         0.471  0.0316 *
Arachnia rubra                              0.468  0.0360 *
Rhodococcus corynebacterioides              0.468  0.0369 *
Irregularibacter muris                      0.467  0.0368 *
Sporosalibacterium tautonense               0.467  0.0306 *
Agrococcus carbonis                         0.465  0.0219 *
Microbacterium halophytorum                 0.465  0.0414 *
Microlunatus phosphovorus NM-1              0.465  0.0418 *
Aliivibrio logei                            0.465  0.0401 *
Salinibacterium hongtaonis                  0.464  0.0442 *
Arthrobacter roseus                         0.463  0.0424 *
Rubripirellula amarantea                    0.462  0.0491 *
Nocardioides houyundeii                     0.460  0.0311 *
Caldicoprobacter faecalis                   0.452  0.0370 *
Desulfobacter postgatei                     0.449  0.0473 *
Rhabdanaerobium thermarum                   0.449  0.0379 *
Naumannella cuiyingiana                     0.448  0.0443 *
Acetonema longum DSM 6540                   0.448  0.0467 *
Terracoccus luteus                          0.448  0.0462 *
Dolosicoccus paucivorans                    0.448  0.0463 *
Georgenia soli                              0.447  0.0441 *
Desulfatibacillum alkenivorans              0.446  0.0379 *
Desulfomonile tiedjei DSM 6799              0.446  0.0414 *
Christensenella hongkongensis               0.445  0.0466 *
Saccharofermentans acetigenes               0.445  0.0490 *
Syntrophomonas palmitatica                  0.443  0.0500 *
Christensenella minuta                      0.438  0.0468 *
Glaciihabitans tibetensis                   0.434  0.0450 *
Caldicoprobacter guelmensis                 0.425  0.0484 *
---
Signif. codes:  0 ‘***’ 0.001 ‘**’ 0.01 ‘*’ 0.05 ‘.’ 0.1 ‘ ’ 1
```

Text S3: Microbial species that showed a statistically significant association in each specific material (Black rock, White rock, HPDE, PE, PET, PVC.A, PLA)/wastewater and their combined groups using a specific R package (indicspecies) to perform Indicator Species Analysis with Norwegian outlet wastewater set-up.

```
Multilevel pattern analysis
 ---------------------------

 Association function: r.g
 Significance level (alpha): 0.05

 Total number of species: 9455
 Selected number of species: 410
 Number of species associated to 1 group: 240
 Number of species associated to 2 groups: 129
 Number of species associated to 3 groups: 34
 Number of species associated to 4 groups: 7
 Number of species associated to 5 groups: 0
 Number of species associated to 6 groups: 0
 Number of species associated to 7 groups: 0

 List of species associated to each combination:

 Group BR  #sps.  12
                            stat p.value
Fulvimonas yonginensis     0.683  0.0274 *
Colwellia piezophila       0.656  0.0150 *
Paracoccus litorisediminis 0.603  0.0273 *
Oceaniferula marina        0.590  0.0230 *
Roseivirga pacifica        0.590  0.0240 *
Rhodopirellula rubra       0.579  0.0483 *
Rubripirellula tenax       0.553  0.0410 *
Paracoccus aestuariivivens 0.553  0.0360 *
Paracoccus limosus         0.549  0.0414 *
Paracoccus yeei            0.546  0.0467 *
Paracoccus denitrificans   0.545  0.0477 *
Paracoccus aminovorans     0.543  0.0488 *

 Group HPDE  #sps.  55
                                            stat p.value
Chryseobacterium scophthalmum              0.874  0.0003 ***
Pseudomonas congelans                      0.847  0.0001 ***
Rhizobium borbori                          0.847  0.0002 ***
Stenotrophomonas humi                      0.840  0.0013 **
Rhizobium paknamense                       0.808  0.0001 ***
Candidimonas humi                          0.808  0.0003 ***
Herbaspirillum autotrophicum               0.800  0.0002 ***
Vibrio aerogenes                           0.798  0.0067 **
Levilactobacillus koreensis JCM 16448      0.798  0.0085 **
Castellaniella defragrans                  0.793  0.0005 ***
Janthinobacterium violaceinigrum           0.746  0.0026 **
Sphingomicrobium astaxanthinifaciens       0.737  0.0027 **
Myroides guanonis                          0.733  0.0043 **
Stenotrophomonas pavanii                   0.723  0.0025 **
Pseudomonas antarctica                     0.720  0.0005 ***
Bordetella trematum                        0.719  0.0007 ***
Stenotrophomonas acidaminiphila            0.714  0.0022 **
Stenotrophomonas nitritireducens           0.712  0.0039 **
Pseudomonas tremae                         0.711  0.0013 **
Flavobacterium gossypii                    0.708  0.0129 *
Pseudomonas ficuserectae                   0.703  0.0021 **
Fulvimarina endophytica                    0.697  0.0030 **
Oligella urethralis                        0.697  0.0027 **
Neorhizobium huautlense                    0.696  0.0032 **
Eikenella corrodens                        0.696  0.0184 *
Oligella ureolytica DSM 18253              0.696  0.0179 *
Roseomonas suffusca                        0.696  0.0173 *
Chryseobacterium arachidis                 0.696  0.0200 *
Chryseobacterium greenlandense             0.696  0.0179 *
Vibrio ezurae                              0.696  0.0195 *
Pseudomonas meridiana                      0.691  0.0021 **
Shewanella xiamenensis                     0.677  0.0045 **
Paracandidimonas soli                      0.665  0.0046 **
Pusillimonas thiosulfatoxidans             0.657  0.0030 **
Janthinobacterium svalbardensis            0.644  0.0152 *
Deefgea rivuli                             0.641  0.0084 **
Pusillimonas caeni                         0.639  0.0057 **
Kocuria uropygioeca                        0.633  0.0244 *
Pseudomonas syringae                       0.629  0.0077 **
Aeromonas salmonicida subsp. pectinolytica 0.620  0.0120 *
Lacticaseibacillus camelliae               0.619  0.0330 *
Aliivibrio sifiae                          0.619  0.0317 *
Rhizobium glycinendophyticum               0.619  0.0101 *
Pusillimonas ginsengisoli                  0.613  0.0098 **
Rhizobium populi                           0.611  0.0138 *
Pseudomonas arsenicoxydans                 0.611  0.0128 *
Tepidimonas taiwanensis                    0.608  0.0227 *
Methylophaga nitratireducenticrescens      0.606  0.0376 *
Stenotrophomonas ginsengisoli              0.603  0.0144 *
Paralcaligenes ginsengisoli                0.596  0.0136 *
Pseudomonas cerasi                         0.584  0.0198 *
Shewanella algae                           0.584  0.0194 *
Xanthomonas axonopodis                     0.571  0.0391 *
Pseudomonas grimontii                      0.552  0.0406 *
Pseudomonas asplenii                       0.547  0.0345 *

 Group PE  #sps.  13
                                 stat p.value
Alkanindiges illinoisensis      0.875  0.0003 ***
Acinetobacter halotolerans      0.836  0.0003 ***
Rhizobium sullae                0.798  0.0013 **
Shewanella aquimarina           0.689  0.0022 **
Mesorhizobium robiniae          0.683  0.0250 *
Microvirgula curvata            0.683  0.0250 *
Pseudomonas borbori             0.629  0.0090 **
Shewanella abyssi               0.603  0.0130 *
Herbaspirillum frisingense      0.601  0.0208 *
Pseudomonas caricapapayae       0.596  0.0180 *
Pseudomonas tolaasii NCPPB 2192 0.581  0.0233 *
Pseudomonas marincola           0.561  0.0301 *
Aeromonas allosaccharophila     0.544  0.0403 *

 Group PET  #sps.  32
                                     stat p.value
Cupriavidus lacunae                 0.683  0.0254 *
Roseomonas aerofrigidensis          0.656  0.0182 *
Constrictibacter antarcticus        0.633  0.0138 *
Novosphingobium colocasiae          0.607  0.0147 *
Methylomonas lenta                  0.601  0.0228 *
Moraxella boevrei                   0.601  0.0217 *
Mesorhizobium mediterraneum         0.598  0.0347 *
Mitsuaria noduli                    0.584  0.0295 *
Psychrobacillus lasiicapitis        0.578  0.0265 *
Flavitalea flava                    0.571  0.0281 *
Sphingomonas koreensis              0.569  0.0322 *
Devosia chinhatensis                0.564  0.0300 *
Comamonas aquatilis                 0.563  0.0290 *
Undibacterium danionis              0.562  0.0428 *
Singulisphaera acidiphila DSM 18658 0.560  0.0339 *
Inhella inkyongensis                0.560  0.0304 *
Legionella spiritensis              0.557  0.0305 *
Pseudoxanthomonas mexicana          0.555  0.0342 *
Rhodocaloribacter litoris           0.554  0.0361 *
Rugosibacter aromaticivorans        0.554  0.0300 *
Anabaena cylindrica PCC 7122        0.553  0.0455 *
Pandoraea terrae                    0.553  0.0445 *
Thiomonas islandica                 0.548  0.0396 *
Pseudoxanthomonas japonensis        0.546  0.0371 *
Nitrosomonas ureae                  0.544  0.0260 *
Caballeronia grimmiae               0.542  0.0455 *
Sphingobium limneticum              0.541  0.0433 *
Herbaspirillum lusitanum            0.538  0.0483 *
Singulisphaera rosea                0.536  0.0365 *
Methylogaea oryzae                  0.535  0.0458 *
Flavobacterium gilvum               0.534  0.0481 *
Ectothiorhodospira magna            0.523  0.0449 *

 Group PLA  #sps.  19
                                    stat p.value
Flavobacterium lindanitolerans     0.798  0.0006 ***
Sphingomonas gotjawalisoli         0.726  0.0016 **
Alishewanella longhuensis          0.706  0.0018 **
Fangia hongkongensis               0.683  0.0267 *
Rheinheimera japonica              0.665  0.0030 **
Rheinheimera baltica               0.658  0.0121 *
Rheinheimera aquimaris             0.630  0.0137 *
Pantoea ananatis                   0.605  0.0147 *
Phaseolibacter flectens ATCC 12775 0.603  0.0209 *
Vibrio halioticoli                 0.601  0.0227 *
Vibrio tapetis subsp. britannicus  0.589  0.0284 *
Sphingobium naphthae               0.585  0.0215 *
Vibrio hispanicus                  0.564  0.0333 *
Rheinheimera marina                0.560  0.0320 *
Pantoea cypripedii                 0.557  0.0304 *
Brenneria alni                     0.554  0.0241 *
Agarivorans litoreus               0.547  0.0489 *
Rheinheimera aestuarii             0.546  0.0455 *
Vibrio penaeicida                  0.546  0.0416 *

 Group PVC.A  #sps.  75
                                                      stat p.value
Brevibacterium ravenspurgense 5401308 = CCUG 53855   0.683  0.0282 *
Microlunatus endophyticus                            0.683  0.0282 *
Schleiferilactobacillus similis DSM 23365 = JCM 2765 0.683  0.0282 *
Campylobacter lanienae NCTC 13004                    0.669  0.0246 *
Pedobacter westerhofensis                            0.660  0.0120 *
Balneicella halophila                                0.660  0.0215 *
Dysgonomonas capnocytophagoides                      0.642  0.0244 *
Parabacteroides goldsteinii DSM 19448 = WAL 12034    0.642  0.0254 *
Pararhodospirillum photometricum DSM 122             0.640  0.0225 *
Polaribacter marinivivus                             0.630  0.0180 *
Persicobacter diffluens                              0.629  0.0180 *
Hydrogenophaga caeni                                 0.625  0.0180 *
Acetobacterium bakii                                 0.624  0.0202 *
Cellulophaga lytica                                  0.622  0.0222 *
Anaerocella delicata                                 0.619  0.0178 *
Alistipes senegalensis JC50                          0.615  0.0180 *
Epilithonimonas bovis DSM 19482                      0.613  0.0303 *
Ancylomarina psychrotolerans                         0.606  0.0187 *
Riemerella anatipestifer                             0.606  0.0229 *
Xenophilus arseniciresistens                         0.604  0.0123 *
Rhodospirillum rubrum ATCC 11170                     0.603  0.0213 *
Abyssisolibacter fermentans                          0.601  0.0239 *
Mediterranea massiliensis                            0.598  0.0333 *
Marininema mesophilum                                0.598  0.0242 *
Gabonia massiliensis                                 0.597  0.0263 *
Sphingobacterium yamdrokense                         0.597  0.0139 *
Cellulomonas carbonis T26                            0.592  0.0357 *
Alistipes finegoldii                                 0.592  0.0263 *
Alistipes shahii                                     0.592  0.0248 *
Alistipes putredinis                                 0.591  0.0256 *
Dysgonomonas gadei ATCC BAA-286                      0.587  0.0212 *
Flavobacterium tegetincola                           0.586  0.0235 *
Acetobacterium psammolithicum                        0.584  0.0173 *
Polaribacter marinaquae                              0.584  0.0259 *
Vallitalea pronyensis                                0.583  0.0305 *
Draconibacterium sediminis                           0.581  0.0215 *
Thiofractor thiocaminus                              0.579  0.0497 *
Alistipes ihumii AP11                                0.579  0.0454 *
Sanguibacter suarezii                                0.579  0.0466 *
Veillonella parvula                                  0.579  0.0466 *
Anaeromicrobium sediminis                            0.577  0.0311 *
Marinifilum albidiflavum                             0.577  0.0339 *
Flavobacterium olei                                  0.576  0.0345 *
Alistipes timonensis JC136                           0.576  0.0271 *
Draconibacterium orientale                           0.575  0.0243 *
Mucilaginibacter daejeonensis                        0.573  0.0305 *
[Clostridium] populeti                               0.572  0.0374 *
Marinifilum fragile CECT 7942                        0.570  0.0497 *
Lutibacter litoralis                                 0.570  0.0326 *
Sphaerochaeta pleomorpha                             0.568  0.0429 *
Acidaminococcus fermentans DSM 20731                 0.568  0.0489 *
Desulfitobacterium chlororespirans                   0.565  0.0401 *
Flavobacterium antarcticum                           0.565  0.0309 *
Phaeocystidibacter marisrubri                        0.565  0.0235 *
Flavobacterium brevivitae                            0.564  0.0490 *
Janibacter hoylei PVAS-1                             0.562  0.0493 *
Saccharicrinis aurantiacus                           0.562  0.0468 *
Williamwhitmania taraxaci                            0.559  0.0368 *
Salibacter halophilus                                0.558  0.0326 *
Azonexus fungiphilus                                 0.555  0.0325 *
Microbacter margulisiae                              0.554  0.0422 *
Vulgatibacter incomptus                              0.553  0.0468 *
Insolitispirillum peregrinum subsp. integrum         0.552  0.0362 *
Desulfofarcimen acetoxidans DSM 771                  0.552  0.0244 *
Mariniphaga sediminis                                0.551  0.0441 *
Desulfitispora elongata                              0.551  0.0472 *
Owenweeksia hongkongensis                            0.550  0.0492 *
[Eubacterium] siraeum                                0.546  0.0388 *
Wandonia haliotis NBRC 105642                        0.546  0.0468 *
Pedobacter mongoliensis                              0.545  0.0209 *
Paenibacillus terreus                                0.543  0.0469 *
Hydromonas duriensis                                 0.541  0.0447 *
Hydrogenophaga soli                                  0.540  0.0485 *
Sulfurospirillum multivorans DSM 12446               0.535  0.0313 *
Dokdonia lutea                                       0.528  0.0430 *

 Group WR  #sps.  2
                      stat p.value
Rickettsia massiliae 0.574  0.0191 *
Aequoribacter fuscus 0.538  0.0412 *

 Group WW  #sps.  32
                                            stat p.value
Listeria innocua                           0.798  0.0088 **
Novipirellula aureliae                     0.753  0.0071 **
Leptolinea tardivitalis                    0.753  0.0088 **
Legionella quateirensis                    0.728  0.0055 **
Nocardioides lianchengensis                0.708  0.0114 *
Rhodococcus rhodochrous                    0.696  0.0185 *
Pelotomaculum thermopropionicum SI         0.677  0.0042 **
Microlunatus sagamiharensis                0.674  0.0016 **
Auraticoccus cholistanensis                0.669  0.0097 **
Nitrosomonas europaea                      0.646  0.0141 *
Peptoniphilus coxii                        0.642  0.0128 *
Microbacterium xylanilyticum               0.619  0.0312 *
Brevibacterium sanguinis                   0.619  0.0305 *
Rhodanobacter humi                         0.619  0.0298 *
Spirosoma jeollabukense                    0.619  0.0305 *
Yersinia nurmii                            0.617  0.0088 **
Anaerofilum agile                          0.613  0.0138 *
Arthrobacter methylotrophus                0.609  0.0030 **
Bacillus alkalitolerans                    0.604  0.0117 *
Microlunatus spumicola                     0.601  0.0046 **
Pengzhenrongella sicca                     0.601  0.0228 *
Microbacterium lacus                       0.599  0.0120 *
Roseomonas aquatica                        0.592  0.0230 *
Serratia proteamaculans                    0.571  0.0208 *
Pseudarthrobacter phenanthrenivorans Sphe3 0.563  0.0300 *
Caloramator australicus RC3                0.562  0.0365 *
Diaminobutyricimonas aerilata              0.557  0.0418 *
Agrococcus terreus                         0.554  0.0444 *
Desulforhopalus singaporensis              0.553  0.0317 *
Serinibacter salmoneus                     0.545  0.0397 *
Desulfobulbus propionicus DSM 2032         0.536  0.0380 *
Fonticella tunisiensis                     0.530  0.0495 *

 Group BR+HPDE  #sps.  2
                          stat p.value
Metabacillus fastidiosus 0.620  0.0278 *
Roseovarius aestuarii    0.541  0.0382 *

 Group BR+PET  #sps.  5
                              stat p.value
Ruegeria meonggei            0.620  0.0155 *
Acinetobacter baylyi         0.579  0.0271 *
Acinetobacter kookii         0.555  0.0496 *
Flavihumibacter cheonanensis 0.544  0.0499 *
Cupriavidus malaysiensis     0.540  0.0461 *

 Group BR+PVC.A  #sps.  1
                          stat p.value
Microbacterium testaceum 0.591  0.0238 *

 Group BR+WW  #sps.  3
                                  stat p.value
Rhodopirellula heiligendammensis 0.687  0.0039 **
Rubripirellula amarantea         0.647  0.0084 **
Nocardioides alpinus             0.601  0.0163 *

 Group HPDE+PE  #sps.  68
                                                 stat p.value
Pseudomonas fildesensis                         0.885  0.0001 ***
Pseudomonas poae                                0.852  0.0001 ***
Pseudomonas qingdaonensis                       0.848  0.0001 ***
Pseudomonas extremaustralis 14-3                0.846  0.0003 ***
Pseudomonas veronii                             0.840  0.0002 ***
Pseudomonas cichorii                            0.832  0.0001 ***
Pseudomonas lurida                              0.829  0.0005 ***
Pseudomonas composti                            0.817  0.0003 ***
Pseudomonas extremorientalis                    0.806  0.0002 ***
Pseudomonas brenneri                            0.800  0.0003 ***
Pseudomonas chlororaphis subsp. aurantiaca      0.799  0.0002 ***
Pseudomonas rhodesiae                           0.791  0.0005 ***
Pseudomonas psychrophila                        0.790  0.0006 ***
Pseudomonas proteolytica                        0.787  0.0011 **
Pseudomonas protegens                           0.777  0.0005 ***
Pigmentiphaga aceris                            0.763  0.0015 **
Pseudomonas trivialis                           0.763  0.0012 **
Pseudomonas fluorescens                         0.752  0.0007 ***
Stenotrophomonas koreensis                      0.734  0.0004 ***
Pseudomonas viridiflava                         0.733  0.0021 **
Pseudomonas marginalis                          0.733  0.0017 **
Pseudomonas chlororaphis                        0.724  0.0011 **
Pseudomonas asturiensis                         0.721  0.0004 ***
Pseudomonas orientalis                          0.710  0.0015 **
Thiomicrospira cyclica ALM1                     0.708  0.0023 **
Pseudomonas tolaasii                            0.706  0.0016 **
Pseudomonas lactis                              0.696  0.0021 **
Pseudomonas costantinii                         0.692  0.0013 **
Pseudomonas plecoglossicida                     0.692  0.0012 **
Pigmentiphaga daeguensis                        0.686  0.0031 **
Pseudomonas weihenstephanensis                  0.684  0.0030 **
Pseudomonas anguilliseptica                     0.682  0.0033 **
Pseudomonas putida                              0.679  0.0013 **
Shewanella pealeana                             0.675  0.0034 **
Pseudomonas panacis                             0.673  0.0025 **
Pseudomonas libanensis                          0.666  0.0047 **
Pseudomonas azotoformans                        0.663  0.0028 **
Pusillimonas noertemannii                       0.663  0.0025 **
Shewanella oneidensis                           0.658  0.0040 **
Shewanella profunda                             0.644  0.0061 **
Pseudomonas agarici                             0.642  0.0035 **
Pseudomonas synxantha                           0.642  0.0073 **
Shewanella japonica                             0.633  0.0074 **
Pseudomonas abietaniphila                       0.623  0.0150 *
Pseudomonas cannabina                           0.619  0.0138 *
Paralcaligenes ureilyticus                      0.617  0.0109 *
[Haemophilus] piscium                           0.613  0.0124 *
Advenella mimigardefordensis DPN7               0.613  0.0112 *
Pseudomonas corrugata                           0.608  0.0128 *
Paenalcaligenes suwonensis                      0.607  0.0109 *
Pseudomonas fragi                               0.602  0.0110 *
Pseudomonas helleri                             0.601  0.0119 *
Pseudomonas gessardii                           0.598  0.0192 *
Shewanella gelidii                              0.594  0.0187 *
Pseudorhizobium marinum                         0.588  0.0185 *
Pseudomonas savastanoi                          0.586  0.0228 *
Pseudomonas aylmerensis                         0.584  0.0257 *
Angulomicrobium tetraedrale                     0.579  0.0354 *
Shewanella amazonensis SB2B                     0.577  0.0231 *
Pseudomonas bohemica                            0.572  0.0244 *
Neisseria chenwenguii                           0.566  0.0307 *
Shewanella seohaensis                           0.566  0.0296 *
Shewanella halifaxensis HAW-EB4                 0.562  0.0278 *
Pseudomonas chlororaphis subsp. aureofaciens    0.561  0.0243 *
Pseudomonas hutmensis                           0.555  0.0376 *
Pseudomonas brassicacearum subsp. neoaurantiaca 0.546  0.0432 *
Pectobacterium betavasculorum                   0.538  0.0490 *
Pseudomonas thivervalensis                      0.537  0.0498 *

 Group HPDE+PET  #sps.  16
                                    stat p.value
Stenotrophomonas tumulicola        0.723  0.0029 **
Noviherbaspirillum aurantiacum     0.681  0.0049 **
Stenotrophomonas pictorum JCM 9942 0.681  0.0045 **
Stenotrophomonas terrae            0.655  0.0077 **
Kushneria indalinina               0.620  0.0246 *
Brevundimonas halotolerans         0.604  0.0126 *
Herminiimonas fonticola            0.595  0.0197 *
Janthinobacterium lividum          0.592  0.0229 *
Herminiimonas saxobsidens          0.591  0.0220 *
Robbsia andropogonis               0.584  0.0239 *
Stenotrophomonas daejeonensis      0.581  0.0283 *
Aquaspirillum arcticum             0.578  0.0298 *
Comamonas guangdongensis           0.576  0.0246 *
Herbaspirillum huttiense           0.566  0.0485 *
Rhizobium azooxidifex              0.557  0.0385 *
Comamonas testosteroni             0.533  0.0446 *

 Group HPDE+PLA  #sps.  2
                             stat p.value
Flavobacterium qiangtangense 0.75  0.0012 **
Flavobacterium kingsejongi   0.64  0.0072 **

 Group HPDE+PVC.A  #sps.  3
                           stat p.value
Alkalicella caledoniensis 0.597  0.0209 *
Paenibacillus contaminans 0.593  0.0190 *
Heliophilum fasciatum     0.541  0.0452 *

 Group HPDE+WW  #sps.  1
                      stat p.value
Advenella faeciporci 0.603  0.0139 *

 Group PE+PLA  #sps.  2
                         stat p.value
Enterovibrio norvegicus 0.570  0.0253 *
Shewanella corallii     0.566  0.0289 *

 Group PE+PVC.A  #sps.  2
                               stat p.value
Nautilia nitratireducens MB-1 0.620  0.0162 *
Neisseria canis               0.617  0.0167 *

 Group PE+WW  #sps.  1
                                     stat p.value
Patulibacter minatonensis DSM 18081 0.644  0.0086 **

 Group PET+PLA  #sps.  10
                                 stat p.value
Pararheinheimera texasensis     0.684  0.0028 **
Rheinheimera pacifica           0.654  0.0065 **
Rheinheimera riviphila          0.638  0.0052 **
Providencia stuartii            0.604  0.0132 *
Brenneria populi Li et al. 2015 0.595  0.0189 *
Rheinheimera nanhaiensis E407-8 0.580  0.0215 *
Rheinheimera perlucida          0.567  0.0266 *
Rheinheimera hassiensis         0.562  0.0284 *
Erythrobacter colymbi           0.541  0.0479 *
Rheinheimera gaetbuli           0.537  0.0499 *

 Group PET+PVC.A  #sps.  1
                      stat p.value
Dechloromonas agitata 0.57  0.0311 *

 Group PET+WW  #sps.  2
                           stat p.value
Friedmanniella luteola    0.655  0.0070 **
Zhihengliuella salsuginis 0.579  0.0208 *

 Group PLA+PVC.A  #sps.  1
                            stat p.value
Leptonema illini DSM 21528 0.591  0.0197 *

 Group PVC.A+WW  #sps.  8
                                             stat p.value
Rhodococcus gannanensis                     0.655  0.0072 **
Clostridium estertheticum subsp. laramiense 0.596  0.0364 *
Auraticoccus monumenti                      0.579  0.0321 *
Tetrasphaera australiensis                  0.577  0.0385 *
Microbacterium profundi                     0.574  0.0360 *
Anaerosinus glycerini                       0.573  0.0425 *
Sporosalibacterium tautonense               0.565  0.0430 *
Desulfoscipio gibsoniae DSM 7213            0.560  0.0469 *

 Group WR+WW  #sps.  1
                        stat p.value
Entomoplasma melaleucae 0.55  0.0375 *

 Group BR+PET+PVC.A  #sps.  1
                                stat p.value
Microbacterium keratanolyticum 0.577  0.0303 *

 Group HPDE+PE+PET  #sps.  20
                                           stat p.value
Janthinobacterium aquaticum               0.690  0.0044 **
Pseudomonas multiresinivorans             0.652  0.0036 **
Aeromonas molluscorum                     0.641  0.0065 **
Duganella levis                           0.624  0.0106 *
Stenotrophomonas maltophilia              0.616  0.0128 *
Janthinobacterium rivuli                  0.589  0.0230 *
Aeromonas jandaei                         0.587  0.0175 *
Pseudomonas silesiensis                   0.581  0.0246 *
Shewanella violacea DSS12                 0.576  0.0255 *
Aeromonas salmonicida subsp. achromogenes 0.574  0.0230 *
Aeromonas salmonicida subsp. smithia      0.568  0.0280 *
Pseudomonas lundensis                     0.564  0.0264 *
Aeromonas encheleia                       0.558  0.0348 *
Janthinobacterium agaricidamnosum         0.554  0.0374 *
Shinella kummerowiae                      0.554  0.0329 *
Lysobacter korlensis                      0.546  0.0441 *
Photobacterium chitinilyticum             0.540  0.0443 *
Pseudomonas lini                          0.539  0.0444 *
Pseudomonas brassicacearum                0.538  0.0411 *
Aeromonas rivuli                          0.537  0.0474 *

 Group HPDE+PE+PLA  #sps.  7
                          stat p.value
Shewanella colwelliana   0.620  0.0110 *
Shewanella pneumatophori 0.614  0.0110 *
Shewanella waksmanii     0.587  0.0202 *
Shewanella putrefaciens  0.580  0.0204 *
Shewanella aestuarii     0.564  0.0295 *
Photobacterium swingsii  0.561  0.0302 *
Pseudomonas peli         0.555  0.0325 *

 Group HPDE+PE+PVC.A  #sps.  2
                       stat p.value
Glaciimonas frigoris  0.560  0.0341 *
Aeromonas salmonicida 0.538  0.0462 *

 Group HPDE+PET+PLA  #sps.  3
                            stat p.value
Sphingobium xenophagum     0.639  0.0068 **
Sphingobium lucknowense F2 0.578  0.0194 *
Alishewanella tabrizica    0.565  0.0258 *

 Group HPDE+PET+WW  #sps.  1
                              stat p.value
Ornithinibacillus salinisoli 0.577  0.0323 *

 Group HPDE+PE+PET+PLA  #sps.  2
                     stat p.value
Moritella viscosa   0.554  0.0348 *
Pseudomonas cedrina 0.532  0.0496 *

 Group HPDE+PE+PET+PVC.A  #sps.  3
                          stat p.value
Caldimonas hydrothermale 0.561  0.0300 *
Acidovorax radicis N35   0.558  0.0344 *
Acidovorax soli          0.545  0.0454 *

 Group HPDE+PE+PET+WW  #sps.  1
                     stat p.value
Pseudomonas migulae 0.555   0.037 *

 Group HPDE+PE+PLA+WW  #sps.  1
                    stat p.value
Shewanella morhuae 0.577  0.0236 *
---
Signif. codes:  0 ‘***’ 0.001 ‘**’ 0.01 ‘*’ 0.05 ‘.’ 0.1 ‘ ’ 1
```

Text S4: Microbial species that showed a statistically significant association in each material group (plastics/rocks) or their combined groups using a specific R package (indicspecies) to perform Indicator Species Analysis with South African inlet and outlet wastewater set-up.

```
 Multilevel pattern analysis
 ---------------------------

 Association function: r.g
 Significance level (alpha): 0.05

 Total number of species: 6469
 Selected number of species: 473
 Number of species associated to 1 group: 473

 List of species associated to each combination:

 Group Plastics  #sps.  101
                                                          stat p.value
Pseudomonas lalkuanensis                                 0.562  0.0068 **
Citrobacter gillenii                                     0.548  0.0043 **
Aeromonas enteropelogenes                                0.536  0.0037 **
Acinetobacter radioresistens                             0.535  0.0067 **
Citrobacter murliniae                                    0.529  0.0018 **
Giesbergeria voronezhensis                               0.512  0.0090 **
Enterobacter sichuanensis                                0.510  0.0069 **
Acinetobacter septicus                                   0.499  0.0077 **
Klebsiella pasteurii                                     0.491  0.0059 **
Tessaracoccus flavescens                                 0.489  0.0084 **
Enterobacter quasiroggenkampii                           0.487  0.0094 **
Comamonas fluminis                                       0.486  0.0122 *
Acinetobacter guillouiae                                 0.485  0.0058 **
Enterobacter chengduensis                                0.481  0.0223 *
Brucella oryzae                                          0.479  0.0155 *
Acinetobacter gyllenbergii                               0.477  0.0170 *
Acinetobacter wuhouensis                                 0.469  0.0067 **
Acinetobacter baumannii                                  0.468  0.0061 **
Delftia deserti                                          0.468  0.0136 *
Shinella pollutisoli                                     0.460  0.0148 *
Enterobacter wuhouensis                                  0.458  0.0070 **
Acinetobacter equi                                       0.456  0.0129 *
Rhodoferax ferrireducens T118                            0.456  0.0196 *
Acinetobacter refrigeratoris                             0.454  0.0121 *
Acinetobacter gandensis                                  0.453  0.0027 **
Roseomonas cervicalis                                    0.452  0.0128 *
Acinetobacter tianfuensis                                0.450  0.0173 *
Rhizobium pseudoryzae                                    0.450  0.0133 *
Giesbergeria sinuosa                                     0.449  0.0181 *
Brevundimonas balnearis                                  0.447  0.0197 *
Phreatobacter cathodiphilus                              0.447  0.0317 *
Brevundimonas mongoliensis                               0.446  0.0235 *
Acinetobacter tandoii                                    0.446  0.0151 *
Acinetobacter baylyi                                     0.444  0.0217 *
Nitratireductor arenosus                                 0.442  0.0160 *
Ramlibacter algicola                                     0.442  0.0260 *
Comamonas suwonensis                                     0.441  0.0196 *
Caulobacter endophyticus                                 0.439  0.0239 *
Comamonas aquatilis                                      0.437  0.0102 *
Acinetobacter junii                                      0.437  0.0169 *
Luteimonas salinisoli                                    0.436  0.0309 *
Paracoccus siganidrum                                    0.436  0.0473 *
Acinetobacter kanungonis                                 0.435  0.0159 *
Acinetobacter shaoyimingii                               0.435  0.0147 *
Neorhizobium petrolearium                                0.430  0.0197 *
Acinetobacter variabilis                                 0.430  0.0144 *
Acinetobacter modestus                                   0.429  0.0242 *
Acinetobacter rongchengensis                             0.427  0.0176 *
Pseudomonas flavescens                                   0.425  0.0468 *
Acinetobacter calcoaceticus                              0.424  0.0244 *
Citrobacter braakii                                      0.420  0.0125 *
Brevundimonas canariensis                                0.418  0.0179 *
Aeromonas veronii                                        0.417  0.0294 *
Acinetobacter indicus                                    0.417  0.0233 *
Rhizobium paknamense                                     0.416  0.0368 *
Acinetobacter bouvetii                                   0.416  0.0192 *
Citrobacter europaeus                                    0.413  0.0044 **
Brevundimonas diminuta ATCC 11568                        0.413  0.0114 *
Acinetobacter lanii                                      0.412  0.0295 *
Vagococcus fluvialis                                     0.409  0.0202 *
Comamonas phosphati                                      0.408  0.0367 *
Brevundimonas terrae                                     0.406  0.0060 **
Acinetobacter johnsonii                                  0.403  0.0191 *
Acinetobacter indicus CIP 110367                         0.401  0.0437 *
Acinetobacter vivianii                                   0.401  0.0405 *
Lampropedia hyalina                                      0.401  0.0462 *
Brevundimonas viscosa                                    0.400  0.0324 *
Agrobacterium leguminum                                  0.399  0.0483 *
Rhizobium borbori                                        0.399  0.0436 *
Acinetobacter albensis                                   0.397  0.0336 *
Pseudocitrobacter vendiensis                             0.397  0.0428 *
Citrobacter freundii ATCC 8090 = MTCC 1658 = NBRC 12681  0.395  0.0083 **
Comamonas koreensis                                      0.395  0.0387 *
Acinetobacter chengduensis                               0.394  0.0267 *
Acinetobacter oryzae                                     0.393  0.0225 *
Acinetobacter pittii DSM 21653                           0.393  0.0392 *
Acinetobacter haemolyticus                               0.392  0.0388 *
Acinetobacter nectaris CIP 110549                        0.391  0.0495 *
Acinetobacter seohaensis                                 0.389  0.0352 *
Acinetobacter bereziniae                                 0.388  0.0348 *
Mycoplana ramosa                                         0.388  0.0351 *
Brevundimonas subvibrioides ATCC 15264                   0.387  0.0222 *
Vagococcus carniphilus                                   0.383  0.0174 *
Comamonas aquatica subsp. rana                           0.383  0.0486 *
Acinetobacter courvalinii                                0.381  0.0447 *
Acinetobacter schindleri                                 0.379  0.0326 *
Brevundimonas diminuta                                   0.376  0.0303 *
Brevundimonas bullata                                    0.374  0.0175 *
Acinetobacter defluvii                                   0.374  0.0144 *
Pseudenterobacter timonensis                             0.372  0.0314 *
Enterococcus hirae ATCC 9790                             0.371  0.0338 *
Acinetobacter oleivorans                                 0.368  0.0495 *
Citrobacter pasteurii                                    0.366  0.0379 *
Clostridium sulfidigenes                                 0.363  0.0369 *
Timonella senegalensis JC301                             0.362  0.0378 *
Chelatococcus daeguensis                                 0.361  0.0397 *
Clostridium subterminale                                 0.348  0.0192 *
Brevundimonas lutea                                      0.336  0.0392 *
Camelimonas fluminis                                     0.326  0.0417 *
Acinetobacter gerneri DSM 14967 = CIP 107464 = MTCC 9824 0.299  0.0443 *
Cloacibacterium haliotis                                 0.299  0.0433 *

 Group Rocks  #sps.  372
                                                   stat p.value
Perlabentimonas gracilis                          0.753  0.0002 ***
Thermosinus carboxydivorans                       0.744  0.0002 ***
Kiritimatiella glycovorans                        0.738  0.0001 ***
Cloacibacillus porcorum                           0.723  0.0001 ***
Maribellus luteus                                 0.722  0.0001 ***
Aminivibrio pyruvatiphilus                        0.714  0.0001 ***
Maribellus maritimus                              0.687  0.0001 ***
Acidaminococcus timonensis                        0.678  0.0001 ***
Solidesulfovibrio magneticus RS-1                 0.676  0.0001 ***
Cloacibacillus evryensis                          0.660  0.0001 ***
Geothrix fermentans                               0.653  0.0002 ***
Desulfomicrobium baculatum DSM 4028               0.648  0.0001 ***
Propionispora vibrioides                          0.645  0.0005 ***
Robertkochia solimangrovi                         0.631  0.0005 ***
Desulfomicrobium norvegicum                       0.629  0.0001 ***
Alistipes finegoldii                              0.627  0.0001 ***
Paludibacterium purpuratum                        0.626  0.0001 ***
Desulfobulbus elongatus                           0.619  0.0002 ***
Zavarzinia compransoris                           0.617  0.0001 ***
Desulfuromonas michiganensis                      0.612  0.0006 ***
Sporomusa malonica                                0.611  0.0004 ***
Desulfobulbus propionicus DSM 2032                0.611  0.0003 ***
Thermanaerovibrio velox                           0.603  0.0001 ***
Solobacterium moorei                              0.581  0.0004 ***
Bacteroides fluxus YIT 12057                      0.579  0.0002 ***
Geobacter pickeringii                             0.577  0.0006 ***
Tardibacter chloracetimidivorans                  0.577  0.0006 ***
Thermanaerovibrio acidaminovorans DSM 6589        0.570  0.0003 ***
Acetonema longum DSM 6540                         0.566  0.0004 ***
Phocaeicola sartorii JCM 16497                    0.561  0.0003 ***
Anaeroarcus burkinensis                           0.561  0.0014 **
Desulfomicrobium hypogeium                        0.559  0.0004 ***
Lentimicrobium saccharophilum                     0.556  0.0002 ***
Alistipes timonensis JC136                        0.555  0.0010 ***
Dongia mobilis                                    0.555  0.0019 **
Meniscus glaucopis                                0.554  0.0011 **
Aminomonas paucivorans DSM 12260                  0.553  0.0010 ***
Veillonella infantium                             0.550  0.0012 **
Megasphaera indica                                0.550  0.0018 **
Anoxybacter fermentans                            0.549  0.0003 ***
Acidaminococcus provencensis                      0.548  0.0030 **
Pusillimonas caeni                                0.547  0.0002 ***
Empedobacter falsenii genomovar 1                 0.545  0.0024 **
Desulfovibrio porci                               0.543  0.0009 ***
Reyranella massiliensis 521                       0.543  0.0005 ***
Sphingomonas prati                                0.542  0.0008 ***
Alistipes shahii                                  0.539  0.0002 ***
Kordiimonas marina                                0.539  0.0009 ***
Lacunisphaera limnophila                          0.539  0.0010 ***
Paracoccus sediminilitoris                        0.539  0.0006 ***
Williamwhitmania taraxaci                         0.536  0.0010 ***
Fusobacterium equinum                             0.534  0.0018 **
Anaerospora hongkongensis                         0.530  0.0015 **
Mucinivorans hirudinis                            0.529  0.0012 **
Eubacterium sulci ATCC 35585                      0.529  0.0030 **
Alistipes shahii WAL 8301                         0.527  0.0010 ***
Maribellus comscasis                              0.526  0.0013 **
Solidesulfovibrio carbinolicus                    0.522  0.0017 **
Verticiella sediminum                             0.518  0.0004 ***
Ancylobacter rudongensis                          0.510  0.0039 **
Sphingobacterium faecale                          0.510  0.0005 ***
Fretibacterium fastidiosum                        0.510  0.0013 **
Desulfonatronum cooperativum                      0.509  0.0011 **
Peptoclostridium acidaminophilum                  0.507  0.0013 **
Horticoccus luteus                                0.504  0.0027 **
Advenella mandrilli                               0.503  0.0030 **
Psychrosinus fermentans                           0.499  0.0015 **
Prolixibacter denitrificans                       0.498  0.0011 **
Lysobacter concretionis                           0.496  0.0171 *
Roseimarinus sediminis                            0.496  0.0006 ***
Paludibacter propionicigenes WB4                  0.494  0.0019 **
Thalassospira australica                          0.491  0.0036 **
Paracandidimonas soli                             0.490  0.0034 **
Desulfobulbus alkaliphilus                        0.488  0.0036 **
Alistipes montrealensis                           0.487  0.0046 **
Anaerosporomusa subterranea                       0.485  0.0074 **
Methylobacillus glycogenes                        0.483  0.0049 **
Desulfoprunum benzoelyticum                       0.480  0.0059 **
Lachnoclostridium pacaense                        0.480  0.0048 **
Streptococcus varani                              0.480  0.0067 **
Desulfovibrio intestinalis                        0.480  0.0017 **
Veillonella magna                                 0.480  0.0014 **
Alkaliflexus imshenetskii                         0.477  0.0051 **
Aquabacter cavernae                               0.477  0.0045 **
Elstera cyanobacteriorum                          0.475  0.0013 **
Castellaniella denitrificans                      0.475  0.0135 *
Pseudostreptobacillus hongkongensis               0.472  0.0070 **
Ralstonia solanacearum                            0.470  0.0086 **
Castellaniella defragrans                         0.470  0.0042 **
Propionivibrio limicola                           0.469  0.0154 *
Cephaloticoccus capnophilus                       0.469  0.0062 **
Geoalkalibacter ferrihydriticus                   0.469  0.0062 **
Legionella hackeliae                              0.469  0.0063 **
Achromobacter aloeverae                           0.468  0.0135 *
Holophaga foetida                                 0.468  0.0018 **
Pusillimonas soli                                 0.468  0.0004 ***
Kaistia hirudinis                                 0.467  0.0023 **
Aquimonas voraii                                  0.467  0.0030 **
Prolixibacter bellariivorans                      0.466  0.0024 **
Propionicimonas paludicola                        0.466  0.0110 *
Prosthecochloris marina                           0.465  0.0010 ***
Castellaniella hirudinis                          0.464  0.0313 *
Kaistia geumhonensis                              0.462  0.0011 **
Eoetvoesia caeni                                  0.460  0.0220 *
Thiovirga sulfuroxydans                           0.460  0.0002 ***
Azospirillum griseum                              0.458  0.0069 **
Opitutus terrae PB90-1                            0.458  0.0064 **
Sunxiuqinia rutila                                0.458  0.0059 **
Aminobacterium colombiense                        0.458  0.0067 **
Pollutimonas nitritireducens                      0.458  0.0034 **
Veillonella atypica                               0.458  0.0047 **
Acetobacteroides hydrogenigenes                   0.456  0.0082 **
Mangrovibacterium marinum                         0.455  0.0020 **
Propionivibrio dicarboxylicus                     0.454  0.0104 *
Dorea phocaeensis                                 0.454  0.0075 **
Lactivibrio alcoholicus                           0.453  0.0036 **
Tahibacter caeni                                  0.452  0.0043 **
Helcococcus sueciensis                            0.451  0.0080 **
Legionella tunisiensis                            0.450  0.0013 **
Labilibacter sediminis                            0.450  0.0071 **
Caenibius tardaugens                              0.450  0.0045 **
Halovulum marinum                                 0.447  0.0049 **
Millionella massiliensis                          0.447  0.0067 **
Rhodopseudomonas palustris                        0.447  0.0049 **
Caenimicrobium hargitense                         0.444  0.0014 **
Fermentimonas caenicola                           0.443  0.0209 *
Pusillimonas thiosulfatoxidans                    0.442  0.0010 ***
Legionella qingyii                                0.442  0.0004 ***
Carboxylicivirga mesophila                        0.442  0.0300 *
Legionella pneumophila subsp. fraseri             0.440  0.0046 **
Reyranella aquatilis                              0.439  0.0025 **
Limisphaera ngatamarikiensis                      0.438  0.0034 **
Achromobacter animicus                            0.437  0.0062 **
Parapusillimonas granuli                          0.436  0.0029 **
Pusillimonas ginsengisoli                         0.435  0.0010 ***
Ampullimonas aquatilis                            0.435  0.0044 **
Legionella drozanskii                             0.435  0.0030 **
Rothia endophytica                                0.435  0.0192 *
Alsobacter metallidurans                          0.434  0.0120 *
Legionella taurinensis                            0.434  0.0049 **
Desulfuromonas svalbardensis                      0.432  0.0138 *
Mangrovibacterium lignilyticum                    0.432  0.0119 *
Syntrophomonas bryantii                           0.432  0.0124 *
Ottowia beijingensis                              0.432  0.0249 *
Flavobacterium celericrescens                     0.430  0.0024 **
Ignavibacterium album JCM 16511                   0.430  0.0068 **
Bifidobacterium adolescentis ATCC 15703           0.430  0.0232 *
Mariniphaga sediminis                             0.428  0.0173 *
Anaeromusa acidaminophila                         0.427  0.0463 *
Pusillimonas minor                                0.426  0.0009 ***
Achromobacter xylosoxidans                        0.426  0.0182 *
Magnetospira thiophila                            0.426  0.0197 *
Pedococcus badiiscoriae                           0.426  0.0210 *
Petrimonas mucosa                                 0.424  0.0108 *
Bacteroides ihuae                                 0.424  0.0088 **
Thermanaeromonas burensis                         0.423  0.0061 **
Flavobacterium cucumis                            0.419  0.0089 **
Chelativorans alearense                           0.418  0.0165 *
Sphingosinicella xenopeptidilytica                0.417  0.0114 *
Azovibrio restrictus                              0.417  0.0043 **
Xinfangfangia soli                                0.416  0.0305 *
Legionella norrlandica                            0.416  0.0008 ***
Bordetella avium                                  0.414  0.0119 *
Oleisolibacter albus                              0.414  0.0081 **
Solidesulfovibrio aerotolerans                    0.413  0.0162 *
Pelistega ratti                                   0.411  0.0136 *
Basilea psittacipulmonis DSM 24701                0.409  0.0211 *
Azonexus hydrophilus DSM 23864                    0.408  0.0114 *
Legionella santicrucis                            0.407  0.0016 **
Bacteroides graminisolvens                        0.407  0.0036 **
Gemmobacter aquaticus                             0.406  0.0341 *
Bacteroides sedimenti                             0.406  0.0163 *
Rossellomorea arthrocnemi                         0.406  0.0152 *
Pusillimonas maritima                             0.405  0.0076 **
Paracandidimonas caeni                            0.405  0.0013 **
Melioribacter roseus P3M-2                        0.403  0.0064 **
Thalassospira lohafexi                            0.403  0.0127 *
Prosthecobacter fluviatilis                       0.403  0.0361 *
Breznakibacter xylanolyticus                      0.402  0.0362 *
Silanimonas mangrovi AK13                         0.401  0.0410 *
Gemmobacter aquatilis                             0.400  0.0163 *
Comamonas serinivorans                            0.400  0.0350 *
Achromobacter dolens                              0.400  0.0070 **
Faecalibacterium duncaniae                        0.399  0.0428 *
Hydrogenoanaerobacterium saccharovorans           0.398  0.0108 *
Legionella massiliensis                           0.398  0.0109 *
Pirellula staleyi DSM 6068                        0.398  0.0130 *
Legionella saoudiensis                            0.397  0.0057 **
Orrella dioscoreae                                0.397  0.0084 **
Oleomonas sagaranensis                            0.397  0.0142 *
Rhizobium tropici CIAT 899                        0.397  0.0244 *
Pseudomonas linyingensis                          0.396  0.0084 **
Alistipes putredinis                              0.394  0.0070 **
Paenalcaligenes niemegkensis                      0.394  0.0198 *
Capillibacterium thermochitinicola                0.393  0.0215 *
Pelistega europaea                                0.393  0.0243 *
Pseudahrensia todarodis                           0.393  0.0226 *
Traorella massiliensis                            0.393  0.0199 *
Desulfobulbus rhabdoformis                        0.393  0.0161 *
Phocaeicola coprophilus                           0.391  0.0130 *
Pygmaiobacter massiliensis                        0.390  0.0176 *
Chryseobacterium reticulitermitis                 0.389  0.0185 *
Sporomusa aerivorans                              0.388  0.0149 *
Legionella sainthelensi                           0.386  0.0207 *
Solidesulfovibrio marrakechensis                  0.386  0.0219 *
Bacteroides uniformis                             0.385  0.0064 **
Agrobacterium rhizogenes                          0.385  0.0479 *
Sporolituus thermophilus DSM 23256                0.385  0.0143 *
Bordetella petrii                                 0.385  0.0180 *
Leucobacter salsicius M1-8                        0.384  0.0264 *
Hydromonas duriensis                              0.384  0.0325 *
Sporomusa acidovorans                             0.383  0.0121 *
Legionella rubrilucens                            0.383  0.0118 *
Achromobacter denitrificans                       0.382  0.0235 *
Advenella kashmirensis                            0.381  0.0330 *
Irregularibacter muris                            0.381  0.0252 *
Tuwongella immobilis                              0.381  0.0129 *
Niveispirillum fermenti                           0.380  0.0295 *
Pseudomonas glareae                               0.379  0.0174 *
Wielerella bovis                                  0.378  0.0209 *
Candidimonas humi                                 0.378  0.0283 *
Alteriqipengyuania halimionae                     0.378  0.0396 *
Amaricoccus macauensis                            0.378  0.0360 *
Anaerotignum lactatifermentans                    0.378  0.0373 *
Anaerovibrio lipolyticus DSM 3074                 0.378  0.0386 *
Bacteroides faecichinchillae JCM 17102            0.378  0.0360 *
Bradyrhizobium canariense                         0.378  0.0372 *
Caecibacteroides pullorum                         0.378  0.0365 *
Duganella alba                                    0.378  0.0381 *
Flavobacterium gillisiae                          0.378  0.0386 *
Hypericibacter terrae                             0.378  0.0403 *
Hyphobacterium vulgare                            0.378  0.0386 *
Kangiella profundi                                0.378  0.0361 *
Legionella steelei                                0.378  0.0365 *
Luteitalea pratensis                              0.378  0.0386 *
Mariniflexile maritimum                           0.378  0.0373 *
Marispirillum indicum                             0.378  0.0392 *
Mogibacterium neglectum                           0.378  0.0372 *
Natronincola histidinovorans                      0.378  0.0376 *
Pukyongia salina                                  0.378  0.0360 *
Rhodococcus rhodnii                               0.378  0.0360 *
Seonamhaeicola marinus                            0.378  0.0361 *
Sphingobium subterraneum                          0.378  0.0389 *
Sphingomonas edaphi                               0.378  0.0389 *
Sunxiuqinia elliptica                             0.378  0.0365 *
Thauera sinica                                    0.378  0.0360 *
[Eubacterium] infirmum                            0.378  0.0410 *
Puteibacter caeruleilacunae                       0.378  0.0062 **
Oceanibaculum pacificum                           0.378  0.0186 *
Curvibacter fontanus                              0.375  0.0464 *
Desulfovibrio simplex                             0.375  0.0377 *
Staphylococcus simulans                           0.374  0.0360 *
Advenella faeciporci                              0.373  0.0083 **
Singulisphaera rosea                              0.371  0.0195 *
Vescimonas coprocola                              0.370  0.0203 *
Afipia carboxidovorans                            0.370  0.0360 *
Gordonia alkaliphila                              0.370  0.0373 *
Tautonia sociabilis                               0.370  0.0361 *
Achromobacter insuavis                            0.368  0.0092 **
Laribacter hongkongensis                          0.368  0.0135 *
Rhodanobacter denitrificans                       0.367  0.0265 *
Paludisphaera soli                                0.366  0.0105 *
Acholeplasma vituli                               0.366  0.0360 *
Desulfobulbus oligotrophicus                      0.366  0.0376 *
Flavobacterium psychrotolerans                    0.363  0.0213 *
Imtechella halotolerans                           0.363  0.0209 *
Pseudoleptotrichia goodfellowii DSM 19756         0.363  0.0214 *
Aliarcobacter trophiarum LMG 25534                0.362  0.0287 *
Hyphomonas polymorpha                             0.361  0.0172 *
Desulfocastanea catecholica                       0.360  0.0386 *
Acetobacterium carbinolicum                       0.356  0.0410 *
Alistipes provencensis                            0.356  0.0410 *
Alterococcus agarolyticus                         0.356  0.0376 *
Apibacter raozihei                                0.356  0.0386 *
Arcobacter vandammei                              0.356  0.0373 *
Bartonella gabonensis                             0.356  0.0360 *
Clostridium thailandense                          0.356  0.0380 *
Coraliomargarita akajimensis                      0.356  0.0379 *
Desulfogranum mediterraneum                       0.356  0.0410 *
Desulfopila inferna                               0.356  0.0386 *
Desulforhopalus singaporensis                     0.356  0.0386 *
Donghicola mangrovi                               0.356  0.0365 *
Eisenbergiella porci                              0.356  0.0373 *
Flavobacterium columnare                          0.356  0.0360 *
Flavobacterium kingsejongi                        0.356  0.0365 *
Hungatella hathewayi                              0.356  0.0386 *
Labrys wisconsinensis                             0.356  0.0386 *
Litorimonas haliclonae                            0.356  0.0410 *
Mangrovitalea sediminis                           0.356  0.0360 *
Methylotenera mobilis JLW8                        0.356  0.0360 *
Motilibacter peucedani                            0.356  0.0386 *
Paraburkholderia xenovorans LB400                 0.356  0.0360 *
Phaeocystidibacter marisrubri                     0.356  0.0376 *
Pseudomonas cichorii                              0.356  0.0373 *
Pseudomonas orientalis                            0.356  0.0373 *
Qipengyuania aerophila                            0.356  0.0360 *
Reyranella graminifolii                           0.356  0.0386 *
Rhodoplanes elegans                               0.356  0.0392 *
Ruminiclostridium hungatei                        0.356  0.0389 *
Sphingomonas piscis                               0.356  0.0378 *
Sunxiuqinia faeciviva                             0.356  0.0386 *
Thermaurantimonas aggregans                       0.356  0.0373 *
Thermodesulfomicrobium thermophilum               0.356  0.0376 *
Thiobacillus sajanensis                           0.356  0.0386 *
Skermanella aerolata                              0.355  0.0481 *
Achromobacter piechaudii                          0.352  0.0154 *
Rhizobium croatiense                              0.352  0.0355 *
Bacteroides cellulosilyticus                      0.352  0.0295 *
Aquihabitans daechungensis                        0.350  0.0373 *
Aromatoleum aromaticum EbN1                       0.350  0.0380 *
Geomobilimonas luticola                           0.350  0.0410 *
Gudongella oleilytica                             0.350  0.0383 *
Lactobacillus amylovorus DSM 20531                0.350  0.0365 *
Marinilabilia rubra                               0.350  0.0364 *
Methylorubrum suomiense                           0.350  0.0380 *
Oryzihumus leptocrescens                          0.350  0.0360 *
Pseudomonas gessardii                             0.350  0.0361 *
Dokdonia aurantiaca                               0.349  0.0373 *
Legionella septentrionalis                        0.349  0.0361 *
Sphingomonas arantia                              0.349  0.0360 *
Macromonas bipunctata                             0.348  0.0171 *
Pusillimonas noertemannii                         0.348  0.0191 *
Legionella rowbothamii                            0.348  0.0361 *
Acidaminococcus intestini                         0.345  0.0277 *
Bacteroides rodentium JCM 16496                   0.345  0.0476 *
Elstera litoralis                                 0.344  0.0252 *
Sphingomonas lenta                                0.344  0.0219 *
Thioclava nitratireducens                         0.344  0.0215 *
Fluviibacter phosphoraccumulans                   0.342  0.0152 *
Alistipes senegalensis JC50                       0.342  0.0123 *
Legionella lytica                                 0.342  0.0239 *
Succiniclasticum ruminis                          0.340  0.0402 *
Anaerovibrio slackiae                             0.333  0.0360 *
Bacteroides ovatus                                0.333  0.0386 *
Rhizorhabdus phycosphaerae                        0.333  0.0360 *
Ercella succinigenes                              0.333  0.0481 *
Novosphingobium naphthalenivorans                 0.333  0.0477 *
Legionella longbeachae                            0.332  0.0170 *
Alysiella filiformis                              0.331  0.0380 *
Isosphaera pallida                                0.331  0.0383 *
Flavobacterium columnare NBRC 100251 = ATCC 23463 0.331  0.0302 *
Aliarcobacter cibarius                            0.330  0.0142 *
Gemmobacter caeni                                 0.330  0.0196 *
Oceanibaculum nanhaiense                          0.329  0.0409 *
Paludisphaera borealis                            0.328  0.0315 *
Phycicoccus duodecadis                            0.328  0.0352 *
Tautonia plasticadhaerens                         0.327  0.0492 *
Desulfovibrio vulgaris                            0.325  0.0352 *
Paracoccus shandongensis                          0.324  0.0493 *
Qipengyuania oceanensis                           0.324  0.0479 *
Aromatoleum bremense                              0.323  0.0421 *
Rivicola pingtungensis                            0.323  0.0365 *
Bacteriovorax stolpii                             0.318  0.0388 *
Brassicibacter thermophilus                       0.318  0.0425 *
Mannheimia ruminalis                              0.318  0.0444 *
Kerstersia similis                                0.314  0.0428 *
Sporomusa paucivorans                             0.314  0.0327 *
Legionella waltersii                              0.313  0.0181 *
Gemmobacter lanyuensis                            0.312  0.0456 *
Sphingosinicella soli                             0.312  0.0495 *
Flavobacterium cheniae                            0.311  0.0375 *
Flavobacterium anseonense                         0.308  0.0360 *
Flavobacterium proteolyticum                      0.302  0.0203 *
Stella humosa                                     0.301  0.0326 *
Aquisphaera giovannonii                           0.297  0.0339 *
Collinsella bouchesdurhonensis                    0.295  0.0462 *
Rhodocaloribacter litoris                         0.295  0.0500 *
Anaerofustis stercorihominis                      0.291  0.0480 *
Flavobacterium buctense                           0.291  0.0482 *
Legionella fallonii                               0.290  0.0497 *
Legionella shakespearei                           0.283  0.0200 *
Flavobacterium dauae                              0.262  0.0392 *
---
Signif. codes:  0 ‘***’ 0.001 ‘**’ 0.01 ‘*’ 0.05 ‘.’ 0.1 ‘ ’ 1
```

Text S5: Microbial species that showed a statistically significant association in each material (Black rock, PET, PVC.A, PLA) or their combined groups using a specific R package (indicspecies) to perform Indicator Species Analysis with South African inlet and outlet wastewater set-up.

```
 Multilevel pattern analysis
 ---------------------------

 Association function: r.g
 Significance level (alpha): 0.05

 Total number of species: 6469
 Selected number of species: 496
 Number of species associated to 1 group: 373
 Number of species associated to 2 groups: 117
 Number of species associated to 3 groups: 6

 List of species associated to each combination:

 Group BR  #sps.  202
                                            stat p.value
Perlabentimonas gracilis                   0.777  0.0001 ***
Maribellus luteus                          0.772  0.0001 ***
Cloacibacillus porcorum                    0.771  0.0001 ***
Kiritimatiella glycovorans                 0.768  0.0001 ***
Aminivibrio pyruvatiphilus                 0.761  0.0001 ***
Acidaminococcus timonensis                 0.719  0.0001 ***
Cloacibacillus evryensis                   0.715  0.0001 ***
Geothrix fermentans                        0.710  0.0002 ***
Solidesulfovibrio magneticus RS-1          0.706  0.0002 ***
Paludibacterium purpuratum                 0.701  0.0002 ***
Thermosinus carboxydivorans                0.682  0.0003 ***
Desulfomicrobium baculatum DSM 4028        0.682  0.0004 ***
Zavarzinia compransoris                    0.678  0.0002 ***
Thermanaerovibrio velox                    0.676  0.0002 ***
Desulfomicrobium norvegicum                0.674  0.0001 ***
Desulfuromonas michiganensis               0.665  0.0005 ***
Geobacter pickeringii                      0.655  0.0021 **
Tardibacter chloracetimidivorans           0.655  0.0018 **
Desulfobulbus elongatus                    0.650  0.0006 ***
Propionispora vibrioides                   0.649  0.0005 ***
Desulfobulbus propionicus DSM 2032         0.648  0.0003 ***
Thermanaerovibrio acidaminovorans DSM 6589 0.640  0.0001 ***
Maribellus maritimus                       0.638  0.0003 ***
Solobacterium moorei                       0.631  0.0002 ***
Lentimicrobium saccharophilum              0.630  0.0001 ***
Alistipes finegoldii                       0.620  0.0004 ***
Kordiimonas marina                         0.617  0.0013 **
Lacunisphaera limnophila                   0.617  0.0016 **
Paracoccus sediminilitoris                 0.617  0.0019 **
Phocaeicola sartorii JCM 16497             0.615  0.0010 ***
Reyranella massiliensis 521                0.613  0.0005 ***
Sporomusa malonica                         0.612  0.0009 ***
Aminomonas paucivorans DSM 12260           0.611  0.0006 ***
Alistipes shahii                           0.606  0.0006 ***
Meniscus glaucopis                         0.605  0.0007 ***
Anoxybacter fermentans                     0.603  0.0007 ***
Robertkochia solimangrovi                  0.598  0.0010 ***
Acetonema longum DSM 6540                  0.595  0.0018 **
Alistipes timonensis JC136                 0.592  0.0018 **
Williamwhitmania taraxaci                  0.590  0.0007 ***
Bacteroides fluxus YIT 12057               0.589  0.0018 **
Fusobacterium equinum                      0.589  0.0042 **
Maribellus comscasis                       0.585  0.0009 ***
Pusillimonas caeni                         0.584  0.0006 ***
Dongia mobilis                             0.583  0.0019 **
Desulfonatronum cooperativum               0.582  0.0008 ***
Desulfomicrobium hypogeium                 0.582  0.0028 **
Anaerospora hongkongensis                  0.582  0.0015 **
Verticiella sediminum                      0.581  0.0010 ***
Sphingobacterium faecale                   0.579  0.0006 ***
Desulfovibrio porci                        0.578  0.0026 **
Veillonella infantium                      0.577  0.0017 **
Mucinivorans hirudinis                     0.576  0.0023 **
Fretibacterium fastidiosum                 0.573  0.0020 **
Prolixibacter denitrificans                0.571  0.0016 **
Roseimarinus sediminis                     0.571  0.0004 ***
Horticoccus luteus                         0.563  0.0059 **
Desulfoprunum benzoelyticum                0.557  0.0120 *
Lachnoclostridium pacaense                 0.557  0.0137 *
Streptococcus varani                       0.557  0.0125 *
Megasphaera indica                         0.555  0.0036 **
Sphingomonas prati                         0.554  0.0042 **
Eubacterium sulci ATCC 35585               0.547  0.0091 **
Thalassospira australica                   0.546  0.0070 **
Cephaloticoccus capnophilus                0.545  0.0108 *
Geoalkalibacter ferrihydriticus            0.545  0.0108 *
Legionella hackeliae                       0.545  0.0117 *
Elstera cyanobacteriorum                   0.543  0.0020 **
Peptoclostridium acidaminophilum           0.542  0.0048 **
Veillonella magna                          0.535  0.0046 **
Advenella mandrilli                        0.535  0.0040 **
Azospirillum griseum                       0.534  0.0125 *
Opitutus terrae PB90-1                     0.534  0.0123 *
Sunxiuqinia rutila                         0.534  0.0120 *
Alistipes montrealensis                    0.533  0.0062 **
Pusillimonas soli                          0.532  0.0006 ***
Thiovirga sulfuroxydans                    0.532  0.0001 ***
Solidesulfovibrio carbinolicus             0.532  0.0058 **
Psychrosinus fermentans                    0.531  0.0049 **
Aquimonas voraii                           0.529  0.0070 **
Aminobacterium colombiense                 0.527  0.0098 **
Lactivibrio alcoholicus                    0.525  0.0047 **
Empedobacter falsenii genomovar 1          0.524  0.0065 **
Anaeroarcus burkinensis                    0.524  0.0057 **
Holophaga foetida                          0.524  0.0027 **
Halovulum marinum                          0.522  0.0109 *
Millionella massiliensis                   0.522  0.0125 *
Rhodopseudomonas palustris                 0.522  0.0109 *
Legionella tunisiensis                     0.520  0.0023 **
Acidaminococcus provencensis               0.519  0.0070 **
Kaistia geumhonensis                       0.518  0.0032 **
Legionella qingyii                         0.516  0.0015 **
Mangrovibacterium marinum                  0.515  0.0075 **
Paracandidimonas soli                      0.513  0.0041 **
Pusillimonas thiosulfatoxidans             0.512  0.0016 **
Pollutimonas nitritireducens               0.511  0.0043 **
Alistipes shahii WAL 8301                  0.509  0.0083 **
Prosthecochloris marina                    0.508  0.0050 **
Ancylobacter rudongensis                   0.508  0.0092 **
Reyranella aquatilis                       0.508  0.0056 **
Castellaniella defragrans                  0.507  0.0083 **
Caenimicrobium hargitense                  0.505  0.0026 **
Kaistia hirudinis                          0.505  0.0095 **
Limisphaera ngatamarikiensis               0.504  0.0056 **
Pusillimonas ginsengisoli                  0.502  0.0016 **
Caenibius tardaugens                       0.502  0.0105 *
Dorea phocaeensis                          0.502  0.0088 **
Parapusillimonas granuli                   0.501  0.0035 **
Methylobacillus glycogenes                 0.500  0.0101 *
Ignavibacterium album JCM 16511            0.499  0.0091 **
Legionella taurinensis                     0.499  0.0074 **
Syntrophomonas bryantii                    0.497  0.0142 *
Legionella pneumophila subsp. fraseri      0.497  0.0084 **
Tahibacter caeni                           0.497  0.0062 **
Prolixibacter bellariivorans               0.496  0.0111 *
Desulfovibrio intestinalis                 0.496  0.0069 **
Ampullimonas aquatilis                     0.494  0.0060 **
Desulfuromonas svalbardensis               0.494  0.0202 *
Mangrovibacterium lignilyticum             0.494  0.0199 *
Castellaniella hirudinis                   0.493  0.0123 *
Pusillimonas minor                         0.493  0.0010 ***
Achromobacter animicus                     0.489  0.0105 *
Legionella norrlandica                     0.489  0.0017 **
Labilibacter sediminis                     0.488  0.0186 *
Aquabacter cavernae                        0.488  0.0130 *
Desulfobulbus alkaliphilus                 0.486  0.0135 *
Flavobacterium celericrescens              0.482  0.0062 **
Legionella santicrucis                     0.480  0.0033 **
Flavobacterium cucumis                     0.480  0.0177 *
Mariniphaga sediminis                      0.480  0.0244 *
Solidesulfovibrio aerotolerans             0.473  0.0237 *
Veillonella atypica                        0.471  0.0150 *
Ralstonia solanacearum                     0.470  0.0203 *
Legionella drozanskii                      0.469  0.0153 *
Paracandidimonas caeni                     0.469  0.0033 **
Castellaniella denitrificans               0.469  0.0191 *
Oleisolibacter albus                       0.469  0.0107 *
Legionella saoudiensis                     0.468  0.0116 *
Magnetospira thiophila                     0.465  0.0430 *
Hydrogenoanaerobacterium saccharovorans    0.464  0.0166 *
Pirellula staleyi DSM 6068                 0.464  0.0156 *
Propionicimonas paludicola                 0.463  0.0204 *
Paludibacter propionicigenes WB4           0.461  0.0213 *
Achromobacter aloeverae                    0.457  0.0259 *
Bacteroides sedimenti                      0.457  0.0283 *
Eoetvoesia caeni                           0.457  0.0264 *
Legionella massiliensis                    0.456  0.0272 *
Azovibrio restrictus                       0.456  0.0204 *
Melioribacter roseus P3M-2                 0.455  0.0154 *
Lysobacter concretionis                    0.455  0.0259 *
Orrella dioscoreae                         0.453  0.0194 *
Oleomonas sagaranensis                     0.453  0.0284 *
Legionella rubrilucens                     0.451  0.0166 *
Achromobacter xylosoxidans                 0.451  0.0420 *
Thermanaeromonas burensis                  0.451  0.0250 *
Acetobacteroides hydrogenigenes            0.451  0.0269 *
Puteibacter caeruleilacunae                0.447  0.0105 *
Bordetella avium                           0.446  0.0227 *
Bacteroides ihuae                          0.446  0.0295 *
Helcococcus sueciensis                     0.446  0.0326 *
Pusillimonas maritima                      0.445  0.0264 *
Rossellomorea arthrocnemi                  0.444  0.0498 *
Alkaliflexus imshenetskii                  0.442  0.0321 *
Alistipes putredinis                       0.440  0.0164 *
Bacteroides graminisolvens                 0.438  0.0147 *
Rothia endophytica                         0.434  0.0362 *
Pseudahrensia todarodis                    0.432  0.0433 *
Anaeromusa acidaminophila                  0.432  0.0426 *
Bifidobacterium adolescentis ATCC 15703    0.431  0.0375 *
Achromobacter denitrificans                0.431  0.0384 *
Sporolituus thermophilus DSM 23256         0.429  0.0316 *
Bordetella petrii                          0.427  0.0398 *
Sphingosinicella xenopeptidilytica         0.427  0.0349 *
Oceanibaculum pacificum                    0.426  0.0346 *
Rhizobium tropici CIAT 899                 0.426  0.0396 *
Advenella faeciporci                       0.425  0.0240 *
Chryseobacterium reticulitermitis          0.425  0.0388 *
Azonexus hydrophilus DSM 23864             0.424  0.0259 *
Chelativorans alearense                    0.423  0.0487 *
Pelistega ratti                            0.422  0.0368 *
Paludisphaera soli                         0.422  0.0253 *
Hyphomonas polymorpha                      0.422  0.0377 *
Leucobacter salsicius M1-8                 0.421  0.0371 *
Bacteroides uniformis                      0.421  0.0406 *
Achromobacter insuavis                     0.418  0.0273 *
Carboxylicivirga mesophila                 0.417  0.0487 *
Candidimonas humi                          0.416  0.0464 *
Achromobacter dolens                       0.415  0.0443 *
Sporomusa acidovorans                      0.414  0.0484 *
Thalassospira lohafexi                     0.414  0.0444 *
Legionella rowbothamii                     0.413  0.0471 *
Laribacter hongkongensis                   0.411  0.0467 *
Hydromonas duriensis                       0.410  0.0495 *
Legionella sainthelensi                    0.408  0.0471 *
Solidesulfovibrio marrakechensis           0.408  0.0443 *
Pusillimonas noertemannii                  0.398  0.0474 *
Legionella longbeachae                     0.395  0.0131 *
Alysiella filiformis                       0.393  0.0354 *
Legionella waltersii                       0.374  0.0292 *
Aliarcobacter cibarius                     0.373  0.0496 *
Flavobacterium proteolyticum               0.361  0.0384 *
Legionella shakespearei                    0.342  0.0329 *

 Group PET  #sps.  39
                                  stat p.value
Acinetobacter beijerinckii       0.653  0.0010 ***
Edwardsiella anguillarum         0.583  0.0010 ***
[Clostridium] viride             0.532  0.0045 **
Propioniciclava soli             0.502  0.0143 *
Dysgonomonas mossii DSM 22836    0.495  0.0114 *
Ramlibacter alkalitolerans       0.490  0.0059 **
Aeromonas schubertii             0.480  0.0134 *
Acinetobacter defluvii           0.480  0.0085 **
Limnohabitans planktonicus II-D5 0.478  0.0158 *
Paraperlucidibaca baekdonensis   0.473  0.0214 *
Aeromonas taiwanensis            0.470  0.0189 *
Intestinimonas timonensis        0.470  0.0152 *
Endothiovibrio diazotrophicus    0.469  0.0283 *
Acinetobacter ihumii             0.454  0.0235 *
Citrobacter braakii              0.451  0.0248 *
Lawsonia intracellularis         0.450  0.0305 *
Lysobacter oligotrophicus        0.450  0.0328 *
Marinobacter oulmenensis         0.450  0.0311 *
Sphingomonas ginkgonis           0.450  0.0315 *
Chitinasiproducens palmae        0.449  0.0208 *
Sporobacter termitidis           0.447  0.0289 *
Peredibacter starrii             0.447  0.0316 *
Acinetobacter baretiae           0.444  0.0462 *
Aquabacterium limnoticum         0.440  0.0300 *
Pantoea dispersa                 0.440  0.0332 *
Tatumella punctata               0.440  0.0308 *
Sphingobium baderi LL03          0.439  0.0343 *
Acinetobacter qingfengensis      0.438  0.0233 *
Atlantibacter hermannii          0.432  0.0485 *
Inediibacterium massiliense      0.432  0.0303 *
Pseudenterobacter timonensis     0.424  0.0339 *
Giesbergeria sinuosa             0.423  0.0458 *
Parabacteroides chongii          0.421  0.0480 *
Clostridium sulfidigenes         0.420  0.0274 *
Citrobacter europaeus            0.420  0.0314 *
Acinetobacter plantarum          0.418  0.0478 *
Phreatobacter cathodiphilus      0.417  0.0500 *
Halothiobacillus neapolitanus    0.415  0.0469 *
Acinetobacter stercoris          0.370  0.0278 *

 Group PLA  #sps.  76
                                       stat p.value
Hydrogenophaga caeni                  0.697  0.0001 ***
Lysobacter chengduensis               0.690  0.0002 ***
Paracoccus siganidrum                 0.676  0.0002 ***
Aromatoleum diolicum                  0.655  0.0017 **
Comamonas terrae                      0.641  0.0003 ***
Novosphingobium endophyticum          0.633  0.0023 **
Paracoccus sulfuroxidans              0.620  0.0003 ***
Sulfurifustis variabilis              0.617  0.0017 **
Oceaniglobus ichthyenteri             0.603  0.0014 **
Agitococcus lubricus                  0.587  0.0030 **
Paracoccus halophilus                 0.576  0.0001 ***
Paracoccus mangrovi                   0.570  0.0023 **
Rhizobium giardinii                   0.568  0.0029 **
Gulbenkiania mobilis                  0.565  0.0038 **
Glutamicibacter protophormiae         0.557  0.0135 *
Ensifer adhaerens                     0.553  0.0033 **
Paracoccus lutimaris                  0.548  0.0039 **
Dechloromonas denitrificans           0.547  0.0035 **
Acidihalobacter prosperus             0.539  0.0078 **
Bartonella ancashensis                0.536  0.0119 *
Brucella papionis                     0.534  0.0112 *
Pseudogemmobacter bohemicus           0.532  0.0033 **
Erythrobacter dokdonensis             0.527  0.0084 **
Chelativorans multitrophicus          0.522  0.0119 *
Lysobacter oculi                      0.519  0.0019 **
Luteimonas yindakuii                  0.519  0.0076 **
Cypionkella sinensis                  0.513  0.0065 **
Xinfangfangia humi                    0.504  0.0034 **
Rhodobacter amnigenus                 0.492  0.0078 **
Vitreoscilla stercoraria              0.492  0.0109 *
Brucella endophytica                  0.490  0.0145 *
Cypionkella collinsensis              0.486  0.0142 *
Paracoccus contaminans                0.483  0.0225 *
Paracoccus zhejiangensis              0.481  0.0161 *
Rhizobium halophytocola               0.481  0.0131 *
Xinfangfangia pollutisoli             0.479  0.0111 *
Pseudorhizobium tarimense             0.477  0.0160 *
Falsigemmobacter faecalis             0.475  0.0130 *
Paracoccus alimentarius               0.473  0.0163 *
Rhizobium indigoferae                 0.472  0.0281 *
Abyssibacter profundi                 0.469  0.0338 *
Hephaestia caeni                      0.466  0.0188 *
Devosia oryziradicis                  0.465  0.0443 *
Sphingobium jiangsuense               0.462  0.0104 *
Luteimonas terricola                  0.462  0.0139 *
Ciceribacter azotifigens              0.462  0.0213 *
Croceibacterium xixiisoli             0.460  0.0254 *
Ciceribacter selenitireducens         0.458  0.0211 *
Pseudogemmobacter hezensis            0.458  0.0135 *
Empedobacter tilapiae                 0.456  0.0166 *
Rhizobium arsenicireducens            0.452  0.0270 *
Eikenella corrodens                   0.446  0.0375 *
Sphingopyxis ginsengisoli             0.446  0.0378 *
Diaphorobacter ruginosibacter         0.445  0.0261 *
Adhaeribacter terreus                 0.444  0.0500 *
Rhizobium skierniewicense Ch11        0.444  0.0202 *
Pseudomonas wadenswilerensis          0.441  0.0357 *
Rhizobium rosettiformans W3           0.441  0.0269 *
Pseudomonas donghuensis               0.440  0.0314 *
Microbacterium saccharophilum         0.440  0.0384 *
Paracoccus koreensis                  0.439  0.0305 *
Roseibium sediminis                   0.436  0.0468 *
Mitsuaria chitinivorans               0.436  0.0372 *
Luteimonas notoginsengisoli           0.435  0.0472 *
Acinetobacter parvus                  0.433  0.0392 *
Novosphingobium colocasiae            0.432  0.0442 *
Tsuneonella dongtanensis              0.432  0.0391 *
Dechlorobacter hydrogenophilus        0.431  0.0349 *
Novosphingobium chloroacetimidivorans 0.431  0.0444 *
Vogesella urethralis                  0.426  0.0016 **
Paracoccus aminophilus                0.425  0.0415 *
Psychrobacter piechaudii              0.419  0.0452 *
Shinella fusca                        0.417  0.0410 *
Mesorhizobium comanense               0.413  0.0486 *
Oryzomicrobium terrae                 0.411  0.0500 *
Ensifer sesbaniae                     0.400  0.0444 *

 Group PVC.A  #sps.  56
                                   stat p.value
Rhizobium pseudoryzae             0.785  0.0001 ***
Caulobacter endophyticus          0.669  0.0001 ***
Roseococcus pinisoli              0.668  0.0001 ***
Limnohabitans australis           0.614  0.0012 **
Camelimonas fluminis              0.613  0.0009 ***
Camelimonas lactis                0.606  0.0008 ***
Desulfosporosinus fructosivorans  0.591  0.0009 ***
Caulobacter segnis                0.581  0.0006 ***
Chelatococcus reniformis          0.576  0.0025 **
Flaviflexus salsibiostraticola    0.543  0.0049 **
Acidovorax wautersii              0.542  0.0039 **
Caulobacter vibrioides            0.535  0.0052 **
Chelatococcus daeguensis          0.530  0.0053 **
Camelimonas abortus               0.527  0.0091 **
Arcanobacterium phocae            0.521  0.0086 **
Ancrocorticia populi              0.511  0.0071 **
Lysobacter xanthus                0.509  0.0100 **
Vagococcus hydrophili             0.506  0.0088 **
Youngiibacter fragilis 232.1      0.506  0.0132 *
Xanthomonas citri pv. malvacearum 0.506  0.0280 *
Comamonas thiooxydans             0.497  0.0073 **
Acidovorax cattleyae              0.494  0.0107 *
Pseudomonas mangrovi              0.489  0.0056 **
Comamonas testosteroni            0.484  0.0096 **
Pararhizobium mangrovi            0.483  0.0133 *
Pelosinus propionicus DSM 13327   0.483  0.0048 **
Methylobacterium durans           0.481  0.0149 *
Melaminivora jejuensis            0.477  0.0032 **
Microvirga subterranea            0.473  0.0175 *
Clostridium disporicum            0.468  0.0345 *
Methylovirgula ligni              0.466  0.0241 *
Acidovorax temperans              0.461  0.0193 *
Changpingibacter yushuensis       0.460  0.0130 *
Flaviflexus huanghaiensis         0.456  0.0105 *
Desulfitispora elongata           0.455  0.0259 *
Pseudomonas insulae               0.454  0.0183 *
Roseomonas cervicalis             0.448  0.0214 *
Merdimmobilis hominis             0.447  0.0338 *
Tessaracoccus coleopterorum       0.444  0.0274 *
Microvirga thermotolerans         0.443  0.0290 *
Caulobacter rhizosphaerae         0.438  0.0184 *
Simplicispira metamorpha          0.437  0.0358 *
Acidovorax antarcticus            0.432  0.0376 *
Anaerosinus glycerini             0.432  0.0084 **
Polaromonas cryoconiti            0.432  0.0343 *
Pseudomonas nitroreducens         0.429  0.0337 *
Acidovorax defluvii               0.429  0.0431 *
Phenylobacterium zucineum HLK1    0.428  0.0415 *
Microlunatus capsulatus           0.425  0.0452 *
Xenophilus arseniciresistens      0.424  0.0415 *
Massilibacillus massiliensis      0.423  0.0300 *
Rhodoferax koreense               0.419  0.0359 *
Acidovorax konjaci                0.417  0.0402 *
Clostridium polynesiense          0.417  0.0475 *
Acidovorax monticola              0.417  0.0463 *
Pseudomonas alcaligenes           0.393  0.0334 *

 Group BR+PET  #sps.  5
                               stat p.value
Anaerosporomusa subterranea   0.493  0.0118 *
Tidjanibacter massiliensis    0.486  0.0123 *
Propionivibrio dicarboxylicus 0.469  0.0167 *
Microbacter margulisiae       0.441  0.0389 *
Limosilactobacillus mucosae   0.428  0.0385 *

 Group BR+PLA  #sps.  14
                                     stat p.value
Gemmobacter aquaticus               0.665  0.0002 ***
Pseudostreptobacillus hongkongensis 0.540  0.0052 **
Alsobacter metallidurans            0.493  0.0134 *
Gemmobacter fontiphilus             0.489  0.0131 *
Novosphingobium bradum              0.481  0.0140 *
Azonexus caeni                      0.470  0.0223 *
Novosphingobium aromaticivorans     0.455  0.0246 *
Novosphingobium arabidopsis         0.447  0.0304 *
Empedobacter brevis                 0.446  0.0340 *
Gemmobacter aquatilis               0.430  0.0328 *
Xinfangfangia soli                  0.427  0.0405 *
Novosphingobium subterraneum        0.416  0.0369 *
Euzebya pacifica                    0.408  0.0431 *
Gemmobacter caeni                   0.396  0.0391 *

 Group BR+PVC.A  #sps.  9
                                 stat p.value
Prosthecobacter dejongeii       0.501  0.0109 *
Prosthecobacter fluviatilis     0.500  0.0100 **
Lacibacterium aquatile          0.457  0.0153 *
Pseudobdellovibrio exovorus JSS 0.455  0.0254 *
Brevifollis gellanilyticus      0.448  0.0213 *
Phenylobacterium kunshanense    0.441  0.0238 *
Prosthecobacter algae           0.435  0.0293 *
Bdellovibrio bacteriovorus      0.420  0.0489 *
Tuwongella immobilis            0.418  0.0452 *

 Group PET+PLA  #sps.  86
                                             stat p.value
Acinetobacter calcoaceticus                 0.603  0.0009 ***
Aeromonas hydrophila                        0.598  0.0013 **
Acinetobacter junii                         0.591  0.0020 **
Acinetobacter radioresistens                0.588  0.0014 **
Comamonas denitrificans                     0.570  0.0025 **
Acinetobacter lanii                         0.565  0.0031 **
Acinetobacter kanungonis                    0.564  0.0028 **
Acinetobacter shaoyimingii                  0.563  0.0032 **
Acinetobacter wuhouensis                    0.554  0.0036 **
Acinetobacter variabilis                    0.551  0.0039 **
Acinetobacter refrigeratoris                0.550  0.0041 **
Acinetobacter tjernbergiae                  0.550  0.0042 **
Acinetobacter rongchengensis                0.548  0.0046 **
Acinetobacter modestus                      0.547  0.0044 **
Acinetobacter bouvetii                      0.547  0.0037 **
Acinetobacter equi                          0.545  0.0053 **
Acinetobacter albensis                      0.543  0.0043 **
Acinetobacter baumannii                     0.542  0.0050 **
Comamonas aquatica subsp. rana              0.541  0.0046 **
Acinetobacter indicus                       0.541  0.0050 **
Acinetobacter courvalinii                   0.541  0.0051 **
Acinetobacter vivianii                      0.540  0.0052 **
Comamonas aquatilis                         0.539  0.0053 **
Acinetobacter tandoii                       0.537  0.0042 **
Acinetobacter haemolyticus                  0.531  0.0064 **
Acinetobacter johnsonii                     0.531  0.0056 **
Comamonas koreensis                         0.530  0.0049 **
Acinetobacter oryzae                        0.528  0.0064 **
Acinetobacter proteolyticus                 0.528  0.0059 **
Comamonas terrigena                         0.527  0.0068 **
Acinetobacter pittii DSM 21653              0.526  0.0050 **
Acinetobacter tianfuensis                   0.524  0.0070 **
Acinetobacter seohaensis                    0.524  0.0063 **
Comamonas jiangduensis                      0.524  0.0074 **
Allohahella antarctica                      0.521  0.0072 **
Comamonas kerstersii                        0.520  0.0059 **
Acinetobacter silvestris                    0.519  0.0071 **
Perlucidibaca piscinae                      0.515  0.0087 **
Pseudacidovorax intermedius                 0.513  0.0096 **
Acinetobacter baylyi                        0.513  0.0065 **
Aeromonas veronii bv. veronii               0.509  0.0094 **
Comamonas aquatica                          0.509  0.0086 **
Acinetobacter lactucae                      0.504  0.0108 *
Acinetobacter soli                          0.498  0.0128 *
Aeromonas lusitana                          0.497  0.0110 *
Mycoplana ramosa                            0.495  0.0088 **
Acinetobacter schindleri                    0.494  0.0117 *
Comamonas nitrativorans                     0.492  0.0149 *
Azonexus fungiphilus                        0.490  0.0143 *
Pseudaeromonas sharmana                     0.490  0.0139 *
Pseudorhodoferax soli                       0.489  0.0122 *
Paracoccus simplex                          0.487  0.0155 *
Acinetobacter bohemicus ANC 3994            0.482  0.0151 *
Cavicella subterranea                       0.482  0.0146 *
Acinetobacter bereziniae                    0.481  0.0149 *
Ciceribacter thiooxidans                    0.480  0.0141 *
Comamonas zonglianii                        0.477  0.0175 *
Acinetobacter oleivorans                    0.475  0.0173 *
Aquirhabdus parva                           0.470  0.0196 *
Acinetobacter halotolerans                  0.468  0.0178 *
Comamonas phosphati                         0.466  0.0219 *
Citrobacter murliniae                       0.465  0.0174 *
Acinetobacter lwoffii                       0.462  0.0227 *
Aeromonas veronii                           0.459  0.0254 *
Acinetobacter guillouiae                    0.456  0.0281 *
Acinetobacter seifertii                     0.456  0.0224 *
Acinetobacter indicus CIP 110367            0.456  0.0259 *
Acinetobacter towneri                       0.454  0.0268 *
Enterococcus aquimarinus                    0.453  0.0253 *
Serpentinimonas barnesii                    0.451  0.0274 *
Acinetobacter venetianus RAG-1 = CIP 110063 0.451  0.0305 *
Aeromonas dhakensis                         0.450  0.0267 *
Acinetobacter brisouii                      0.450  0.0251 *
Thalassomonas viridans                      0.445  0.0330 *
Pseudaeromonas pectinilytica                0.445  0.0261 *
Rhizobium sullae                            0.444  0.0358 *
Rhizobium borbori                           0.444  0.0336 *
Acinetobacter apis                          0.443  0.0332 *
Acinetobacter nectaris CIP 110549           0.440  0.0354 *
Luteimonas granuli                          0.435  0.0342 *
Novosphingobium hassiacum                   0.434  0.0408 *
Pseudorhizobium marinum                     0.432  0.0360 *
Aeromonas enteropelogenes                   0.426  0.0457 *
Enterobacter wuhouensis                     0.424  0.0380 *
Pseudomonas argentinensis                   0.423  0.0440 *
Acinetobacter wanghuae                      0.416  0.0428 *

 Group PET+PVC.A  #sps.  2
                          stat p.value
Brevundimonas aurantiaca 0.473  0.0198 *
Brevundimonas viscosa    0.424  0.0380 *

 Group PLA+PVC.A  #sps.  1
                      stat p.value
Diaphorobacter caeni 0.498   0.008 **

 Group BR+PET+PLA  #sps.  4
                               stat p.value
Propionivibrio limicola       0.528  0.0053 **
Macellibacteroides fermentans 0.524  0.0059 **
Parabacteroides chartae       0.502  0.0095 **
Sphingomonas laterariae       0.417  0.0485 *

 Group PET+PLA+PVC.A  #sps.  2
                          stat p.value
Pseudomonas lalkuanensis 0.446  0.0286 *
Citrobacter gillenii     0.425  0.0409 *
---
Signif. codes:  0 ‘***’ 0.001 ‘**’ 0.01 ‘*’ 0.05 ‘.’ 0.1 ‘ ’ 1
```

Table S1: AMR genes detected using staramr tool

| **Gene** | **Predicted Phenotype** | **material** | **source** |
| --- | --- | --- | --- |
| mph(E) | erythromycin, azithromycin | PLA | Inlet |
| msr(E) | erythromycin, azithromycin | PLA | Inlet |
| tet(39) | tetracycline | PLA | Inlet |
| sul1 | sulfisoxazole | PET | Inlet |
| aph(6)-Id | kanamycin | PE | Inlet |
| sul2 | sulfisoxazole | PE | Inlet |
| aac(3)-Ib | gentamicin | PLA | Outlet |
| blaBEL-1 | ampicillin, amoxicillin/clavulanic acid, cefoxitin, ceftriaxone | PLA | Outlet |
| blaOXA-539 | ampicillin | PLA | Outlet |
| qacE | unknown[qacE_1_X68232] | PLA | Outlet |
| sul1 | sulfisoxazole | PLA | Outlet |
| blaOXA-58 | ampicillin, meropenem | PET | Outlet |
| blaOXA-58 | ampicillin, meropenem | PET | Outlet |
| sul1 | sulfisoxazole | PE | Outlet |

Table S2: Mobile genetic elements detected on chromosome using mobsuite tool.

| **molecule_type** | **contig_id** | **mge_type** | **mge_subtype** | **mge_length** | **material** | **source** |
| --- | --- | --- | --- | --- | --- | --- |
| chromosome | contig_5 | ISPsy42 | Tn3 | 5667 | PLA | Inlet |
| chromosome | contig_5 | ISPsy42 | Tn3 | 5667 | PLA | Inlet |
| chromosome | contig_365 | IS6100 | IS6 | 5466 | PET | Inlet |
| chromosome | contig_365 | IS6100 | IS6 | 5466 | PET | Inlet |
| chromosome | contig_75 | IS401 | IS3 | 1316 | PET | Inlet |
| chromosome | contig_10 | ISAlw1 | IS5 | 1039 | BR | Outlet |
| chromosome | contig_136 | ISPpu12 | ISL3 | 3372 | BR | Outlet |
| chromosome | contig_262 | ISAba12 | IS5 | 1039 | BR | Outlet |
| chromosome | contig_3 | ISAba21 | IS3 | 1274 | BR | Outlet |
| chromosome | contig_544 | ISAba14 | IS3 | 1283 | BR | Outlet |
| chromosome | contig_111 | ISSod25 | IS91 | 2313 | PLA | Outlet |
| chromosome | contig_1144 | ISAba125 | IS30 | 2175 | PLA | Outlet |
| chromosome | contig_121 | ISPa22 | IS1182 | 1669 | PLA | Outlet |
| chromosome | contig_126 | ISPa22 | IS1182 | 1669 | PLA | Outlet |
| chromosome | contig_129 | ISCARN66 | IS5 | 1200 | PLA | Outlet |
| chromosome | contig_159 | IS1474 | IS21 | 2595 | PLA | Outlet |
| chromosome | contig_161 | ISPsme1 | IS30 | 1066 | PLA | Outlet |
| chromosome | contig_17 | ISVapa4 | IS21 | 2605 | PLA | Outlet |
| chromosome | contig_190 | ISPpu18 | IS5 | 1192 | PLA | Outlet |
| chromosome | contig_192 | ISSpu5 | IS21 | 2481 | PLA | Outlet |
| chromosome | contig_20 | ISAba21 | IS3 | 1274 | PLA | Outlet |
| chromosome | contig_223 | ISPa1635 | IS4 | 1637 | PLA | Outlet |
| chromosome | contig_223 | ISButh6 | IS5 | 1331 | PLA | Outlet |
| chromosome | contig_223 | ISButh6 | IS5 | 1331 | PLA | Outlet |
| chromosome | contig_224 | ISPpu18 | IS5 | 1192 | PLA | Outlet |
| chromosome | contig_225 | ISCARN66 | IS5 | 1200 | PLA | Outlet |
| chromosome | contig_269 | ISPpu18 | IS5 | 1192 | PLA | Outlet |
| chromosome | contig_349 | ISSod25 | IS91 | 2313 | PLA | Outlet |
| chromosome | contig_387 | IS1474 | IS21 | 2595 | PLA | Outlet |
| chromosome | contig_4 | ISPa1635 | IS4 | 1637 | PLA | Outlet |
| chromosome | contig_4 | ISRme10 | IS30 | 1113 | PLA | Outlet |
| chromosome | contig_404 | ISApr11 | IS1380 | 1671 | PLA | Outlet |
| chromosome | contig_405 | ISApr11 | IS1380 | 1671 | PLA | Outlet |
| chromosome | contig_48 | ISRme10 | IS30 | 1113 | PLA | Outlet |
| chromosome | contig_523 | ISRme10 | IS30 | 1113 | PLA | Outlet |
| chromosome | contig_61 | ISAba125 | IS30 | 2175 | PLA | Outlet |
| chromosome | contig_64 | IS1474 | IS21 | 2595 | PLA | Outlet |
| chromosome | contig_735 | ISAba12 | IS5 | 1039 | PLA | Outlet |
| chromosome | contig_75 | ISPpu12 | ISL3 | 3372 | PLA | Outlet |
| chromosome | contig_75 | ISVapa4 | IS21 | 2605 | PLA | Outlet |
| chromosome | contig_78 | ISButh6 | IS5 | 1331 | PLA | Outlet |
| chromosome | contig_78 | ISPpu12 | ISL3 | 3372 | PLA | Outlet |
| chromosome | contig_81 | ISRme10 | IS30 | 1113 | PLA | Outlet |
| chromosome | contig_82 | IS1474 | IS21 | 2595 | PLA | Outlet |
| chromosome | contig_82 | ISSpu5 | IS21 | 2481 | PLA | Outlet |
| chromosome | contig_844 | ISStma11 | ISL3 | 4426 | PLA | Outlet |
| chromosome | contig_85 | ISVapa4 | IS21 | 2605 | PLA | Outlet |
| chromosome | contig_879 | ISSba6 | IS4 | 1420 | PLA | Outlet |
| chromosome | contig_883 | ISCARN66 | IS5 | 1200 | PLA | Outlet |
| chromosome | contig_92 | ISRme10 | IS30 | 1113 | PLA | Outlet |
| chromosome | contig_92 | ISCARN66 | IS5 | 1200 | PLA | Outlet |
| chromosome | contig_1283 | ISAba12 | IS5 | 1039 | PET | Outlet |
| chromosome | contig_1290 | ISPpu18 | IS5 | 1192 | PET | Outlet |
| chromosome | contig_1458 | ISRme10 | IS30 | 1113 | PET | Outlet |
| chromosome | contig_1505 | ISPa22 | IS1182 | 1669 | PET | Outlet |
| chromosome | contig_165 | ISAba125 | IS30 | 2175 | PET | Outlet |
| chromosome | contig_167 | ISAba125 | IS30 | 2175 | PET | Outlet |
| chromosome | contig_168 | ISAba12 | IS5 | 1039 | PET | Outlet |
| chromosome | contig_168 | ISAba125 | IS30 | 2175 | PET | Outlet |
| chromosome | contig_173 | ISAlw1 | IS5 | 1039 | PET | Outlet |
| chromosome | contig_174 | ISAba17 | IS66 | 2491 | PET | Outlet |
| chromosome | contig_175 | ISAba14 | IS3 | 1283 | PET | Outlet |
| chromosome | contig_175 | ISAba125 | IS30 | 2175 | PET | Outlet |
| chromosome | contig_183 | ISAba21 | IS3 | 1274 | PET | Outlet |
| chromosome | contig_183 | ISAba21 | IS3 | 1274 | PET | Outlet |
| chromosome | contig_183 | ISAba21 | IS3 | 1274 | PET | Outlet |
| chromosome | contig_19 | ISAba11 | IS701 | 1101 | PET | Outlet |
| chromosome | contig_19 | ISAba14 | IS3 | 1283 | PET | Outlet |
| chromosome | contig_191 | ISSpe2 | IS110 | 1366 | PET | Outlet |
| chromosome | contig_212 | ISAba12 | IS5 | 1039 | PET | Outlet |
| chromosome | contig_226 | IS1474 | IS21 | 2595 | PET | Outlet |
| chromosome | contig_227 | ISRme10 | IS30 | 1113 | PET | Outlet |
| chromosome | contig_271 | ISSpe2 | IS110 | 1366 | PET | Outlet |
| chromosome | contig_286 | ISSpe2 | IS110 | 1366 | PET | Outlet |
| chromosome | contig_300 | ISStma11 | ISL3 | 4426 | PET | Outlet |
| chromosome | contig_300 | ISStma11 | ISL3 | 4426 | PET | Outlet |
| chromosome | contig_350 | ISSde6 | IS3 | 1235 | PET | Outlet |
| chromosome | contig_350 | ISSod25 | IS91 | 2313 | PET | Outlet |
| chromosome | contig_37 | ISRme10 | IS30 | 1113 | PET | Outlet |
| chromosome | contig_37 | ISPpu12 | ISL3 | 3372 | PET | Outlet |
| chromosome | contig_37 | ISButh6 | IS5 | 1331 | PET | Outlet |
| chromosome | contig_37 | ISPst2 | ISL3 | 2985 | PET | Outlet |
| chromosome | contig_37 | ISButh6 | IS5 | 1331 | PET | Outlet |
| chromosome | contig_40 | ISAba12 | IS5 | 1039 | PET | Outlet |
| chromosome | contig_40 | ISAlw1 | IS5 | 1039 | PET | Outlet |
| chromosome | contig_406 | ISAba125 | IS30 | 2175 | PET | Outlet |
| chromosome | contig_406 | ISAba125 | IS30 | 2175 | PET | Outlet |
| chromosome | contig_414 | ISSba6 | IS4 | 1420 | PET | Outlet |
| chromosome | contig_457 | ISAba17 | IS66 | 2491 | PET | Outlet |
| chromosome | contig_498 | ISAeme12 | IS4 | 1426 | PET | Outlet |
| chromosome | contig_498 | ISKpn26 | IS5 | 1197 | PET | Outlet |
| chromosome | contig_56 | ISAs31 | IS3 | 1322 | PET | Outlet |
| chromosome | contig_6 | ISAba24 | IS66 | 2421 | PET | Outlet |
| chromosome | contig_695 | ISApr11 | IS1380 | 1671 | PET | Outlet |
| chromosome | contig_856 | ISSde6 | IS3 | 1235 | PET | Outlet |
| chromosome | contig_856 | ISSde6 | IS3 | 1235 | PET | Outlet |
| chromosome | contig_90 | ISPa1635 | IS4 | 1637 | PET | Outlet |
| chromosome | contig_906 | ISKpn31 | ISAs1 | 1441 | PET | Outlet |
| chromosome | contig_925 | ISAba21 | IS3 | 1274 | PET | Outlet |
| chromosome | contig_96 | ISRme10 | IS30 | 1113 | PET | Outlet |
| chromosome | contig_96 | ISPst3 | IS21 | 2606 | PET | Outlet |
| chromosome | contig_96 | ISPst3 | IS21 | 2606 | PET | Outlet |
| chromosome | contig_962 | ISCARN66 | IS5 | 1200 | PET | Outlet |
| chromosome | contig_103 | ISKpn26 | IS5 | 1197 | PE | Outlet |
| chromosome | contig_138 | ISSpe2 | IS110 | 1366 | PE | Outlet |
| chromosome | contig_172 | ISVapa4 | IS21 | 2605 | PE | Outlet |
| chromosome | contig_172 | ISPa22 | IS1182 | 1669 | PE | Outlet |
| chromosome | contig_173 | IS1474 | IS21 | 2595 | PE | Outlet |
| chromosome | contig_203 | ISAs17 | IS3 | 1333 | PE | Outlet |
| chromosome | contig_282 | ISApr11 | IS1380 | 1671 | PE | Outlet |
| chromosome | contig_317 | ISPpu18 | IS5 | 1192 | PE | Outlet |
| chromosome | contig_324 | ISRme10 | IS30 | 1113 | PE | Outlet |
| chromosome | contig_425 | ISPpu12 | ISL3 | 3372 | PE | Outlet |
| chromosome | contig_44 | ISSpe2 | IS110 | 1366 | PE | Outlet |
| chromosome | contig_466 | ISSde6 | IS3 | 1235 | PE | Outlet |
| chromosome | contig_474 | ISVapa4 | IS21 | 2605 | PE | Outlet |
| chromosome | contig_59 | ISSba6 | IS4 | 1420 | PE | Outlet |
| chromosome | contig_601 | ISPpu18 | IS5 | 1192 | PE | Outlet |
| chromosome | contig_677 | ISAs31 | IS3 | 1322 | PE | Outlet |

Table S2: Mobile genetic elements detected on plasmid using mobsuite tool.

| **molecule_type** | **contig_id** | **mge_type** | **mge_subtype** | **mge_length** | **material** | **source** |
| --- | --- | --- | --- | --- | --- | --- |
| plasmid | contig_1470 | ISPa38 | Tn3 | 3400 | PET | Outlet |
| plasmid | contig_573 | ISAba11 | IS701 | 1101 | PET | Outlet |
